# Blind sparse deconvolution for inferring spike trains from fluorescence recordings

**DOI:** 10.1101/156364

**Authors:** Jérôme Tubiana, Sébastien Wolf, Georges Debregeas

## Abstract

The parallel developments of genetically-encoded calcium indicators and fast fluorescence imaging techniques makes it possible to simultaneously record neural activity of extended neuronal populations *in vivo*, opening a new arena for systems neuroscience. To fully harness the potential of functional imaging, one needs to infer the sequence of action potentials from fluorescence time traces. Here we build on recently proposed computational approaches to develop a blind sparse deconvolution algorithm (BSD), which we motivate by a theoretical analysis. We demonstrate that this method outperforms existing sparse deconvolution algorithms in terms of robustness, speed and/or accuracy on both synthetic and real fluorescence data. Furthermore, we provide solutions for the practical problems of thresholding and determination of the rise and decay time constants. We provide theoretical bounds on the performance of the algorithm in terms of precision-recall and temporal accuracy. Finally, we extend the computational framework to support temporal superresolution whose performance is established on real data.

## Introduction

In the last two decades, functional calcium imaging has emerged as a popular method for recording brain activity *in-vivo*. This technique relies on calcium sensors, either synthetic or genetically expressed, that are designed to optically report the transient rise in intra-cellular calcium concentration that accompany spiking events. Calcium imaging offers several assets in comparison with standard electrophysiology methods: it is non-invasive, it allows monitoring extended neuronal networks (up to a few tens of thousands of units), and it can be combined with genetic methods in order to target specific neuronal populations.

The main limitation of calcium imaging is that it only provides a proxy measure of the neuronal activity. The kinetics of the calcium/reporter complexation being relatively slow, the spike-evoked fluorescence transients last much longer (0.1-1s) than the action potential itself (*<*5ms). As the fluorescence signal is generally noisy and/or weakly sampled, its interpretation heavily relies on deconvolution methods to infer approximated spike trains. With the rapid increase in data-throughput offered by current fast imaging techniques, these methods need to be fast and unsupervised, as any manual check of the produced inference signals would prove impractical.

Standard inference methods are based on a generative model, which describes the relationship between a spike train and the resulting fluorescence time trace. It reads:

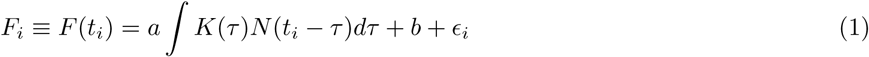

where *t*_*i*_ = *i*Δ*t* is the time of measurement, 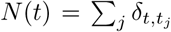 denotes the spike train, *b* is the baseline fluorescence (spikeless signal), and *ϵ*_*i*_ is a discrete gaussian white noise: *< ϵ*_*i*_ *>*= 0, *< ϵ*_*i*_*ϵ*_*j*_ *>*= *σ*^2^*δ*_*i,j*_. The convolution kernel *K*(*t*), which reflects the complexation kinetics, is of the form:

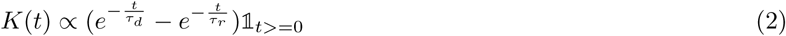

where the rise and decay time constants *τ*_*r*_, *τ*_*d*_ – typically in the range of 10-100ms and 50-1000ms, respectively – mostly depend on the calcium indicator but can also vary with the targeted neuron. In the following, we normalize *K* such that max(*K*) = 1, hence each spike produces a transient of maximum height *a*. The signal to noise ratio (SNR) is thus defined as 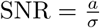. The noise stems from fluctuations of intra-cellular chemical concentration, light source and detector noise, incorrect baseline estimation, and other modeling errors. Because the SNR is often small in practice (*∈* [2, 10]) simple inference methods such as naive linear deconvolution and Wiener filtering are inadequate. In the last decade, numerous alternative deconvolution algorithms have been proposed [1–15]; among them, a powerful family of algorithms is based on non-negative sparse deconvolution [5, 13, 14]. In short, it consists in solving the inverse problem (equation (1)) using the *a priori* knowledge that the spikes are sparse and non-negative. This framework, introduced by Vogelstein *et al.* in [5], was shown to efficiently recover spike trains from fluorescence signals. However, its performance is strongly dependent on the algorithm’s hyperparameters, namely the sparsity prior *λ* and the rise and decay time constants *τ*_*r*_, *τ*_*d*_. Despite extensive efforts for automatically adjusting these parameters [13, 14], progress are still needed to achieve the adequate robustness of the inference [16]. Another drawbacks of such algorithms is that the interpretation of the output can be challenging due to a paucity of theoretical understanding of its expected performance. First, the inferred signal is a continuous time series, which thus needs to be thresholded to yield a discrete set of spikes. Current methods remain elusive regarding how the threshold value controls the precision-recall of spike detection. Second, the temporal accuracy of the ouptut signal remains elusive: the probability that a given spike be inferred at the wrong time bin - in advance or delayed with respect to the true spike - is unkown. Conversely, one may expect that for high SNR configurations, the temporal resolution of the inferred signal may exceed the fluorescence sampling rate.

In the context of our laser-sheet calcium imaging experiments on zebrafish larvae, we built on the fast-oopsi algorithm to develop a so-called *Blind Sparse Deconvolution (BSD)* algorithm. This implementation features automatic estimation of the hyperparameters, enhanced speed and similar-to-better reconstruction performances and super-resolution capabilities. It is benchmarked on both synthetic and real data. We additionally provide thresholding guidelines and theoretical bounds on its performance, in terms of precision-recall and temporal accuracy, for large range of realistic experimental conditions. The program is available at <u>https://github.com/jertubiana/BSD</u>.

The article is organized as follows. In section 1, we present the principles of non-negative sparse deconvolution methods. In section 2, we focus on the determination of the sparsity prior *λ*; we compare our method to existing solutions and compare performances on synthetic data. In section 3, we compute the spike detection theoretical limits in terms of precision-recall and temporal resolution. In section 4, we focus on the automatic determination of the parameters of the generative model, in particular the time-constants of the convolution kernel *K*. In section 5, we extend BSD to support super-resolution, and characterize the resulting gain in temporal resolution on synthetic data. Finally, in section 6, we test our algorithm on real data, (i) fluorescence recordings in mice cortex with electrophysiology ground truth [17, 18], and (ii) whole-brain recordings of larval zebrafish [19, 20].

## I. NON-NEGATIVE SPARSE DECONVOLUTION

In this first section, we review the existing deconvolution approaches for inferring *N* (*t*). We rewrite Eqn. 1 as:

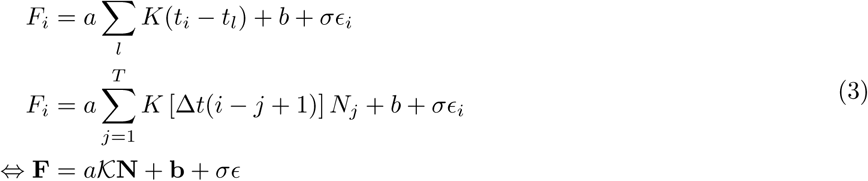

where *i ∈* [1, *T*] is the time frame index, 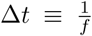 is the sampling interval, *K* is the convolution matrix *K*_*ij*_ = *K* [Δ*t*(*i − j* + 1)] and 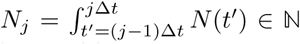 is the number of spikes in the time interval [(*j −* 1)Δ*t, j*Δ*t*] [29]. Note that the first and second lines are not equivalent; in doing so, we assume that:

- The boundary condition *N* (*t*) = 0, *∀ t <* 0 holds. It is mostly true in typical recordings that start during inactive periods but this simplification can be easily relaxed. [30]
- We can approximate *K*(*t*_*i*_ *− t*_*l*_) = *K*(*i*Δ*t − t*_*l*_) as *K* [*i*Δ*t −*(*j*_*l*_ *−* 1)Δ*t*] where 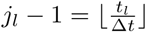. This discretization error is negligible when Δ*t* is small, yet it ensures that the matrix *K* is translation invariant, *i.e. K* _*ij*_ = *φ*(*i − j*) [31].

Following 3, a naive estimate for *N* is:

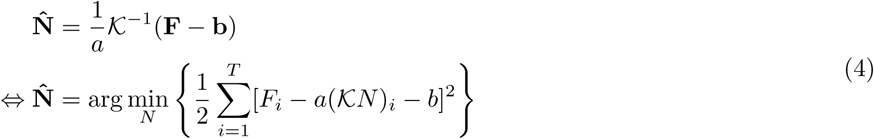

As shown in Fig. 1a, this approach fails to recover any spike at typical noise level *SNR* = 2.5. To understand this failure, one may reason in the continuous framework for which Eqn. 4 writes 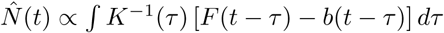 Here, the inverse convolution kernel *K*^−1^ is proportional to 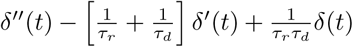 thus the naive deconvo-lution reads:

**FIG. 1:**
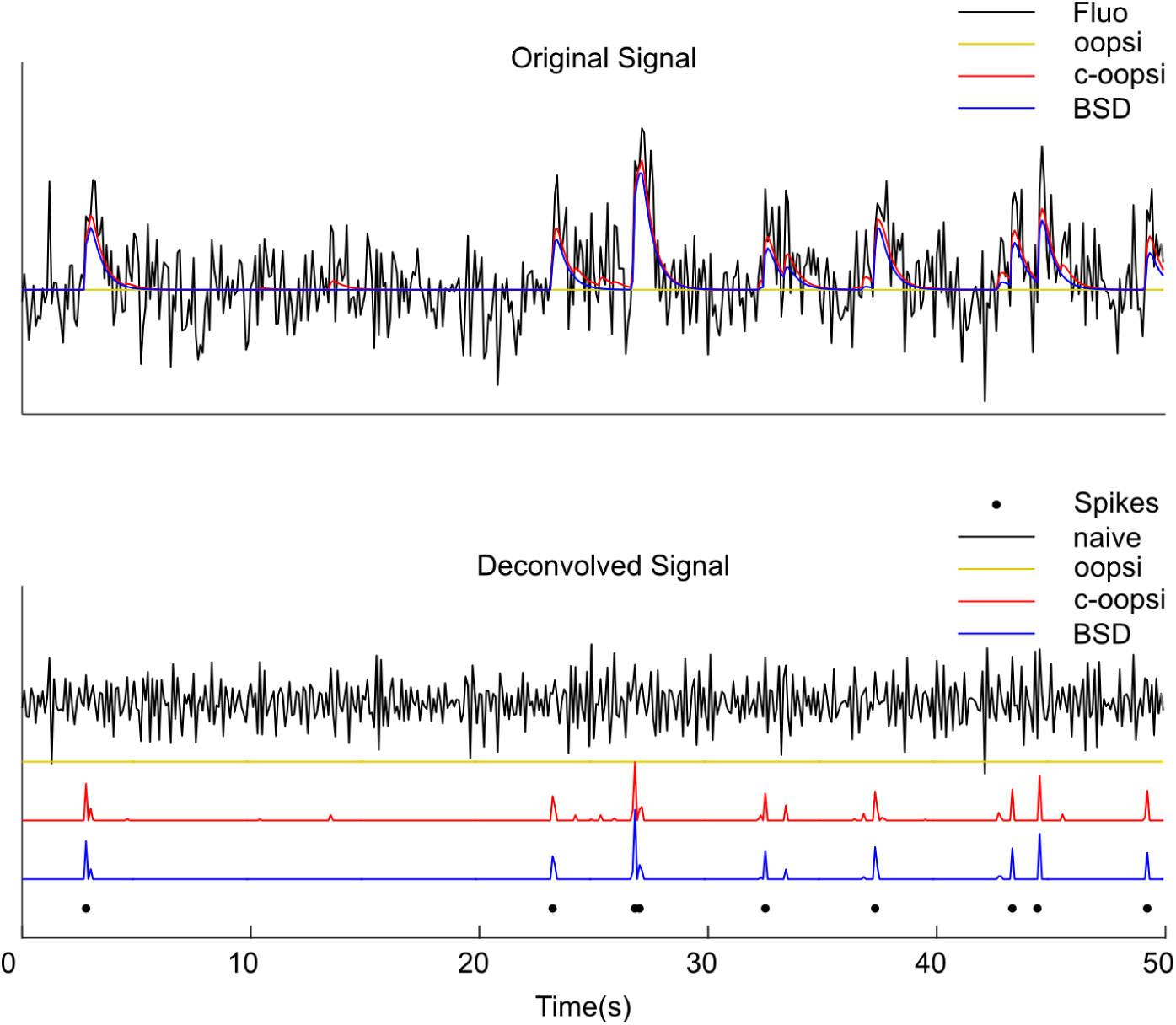
Example of inference results.

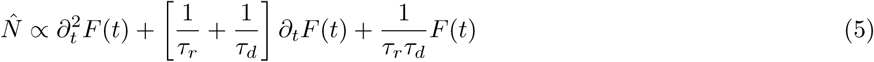

A naive estimator of the signal involves computing the derivatives of the original signal, and is thus extremely sensitive to high frequency noise. The reasoning is similar for discrete signals, which involve discrete time derivatives. An intuitive solution to mitigate this issue consists in filtering out the high frequency component before carrying out the deconvolution, as is the basis of the Wiener deconvolution method. Vogelstein et al. showed that it also performs poorly because it smoothes out the fast rise of the fluorescence signal at spikes. In contrast, non-negative sparse deconvolution estimators achieve both filtering of the noise while preserving the high-frequency signal. They are given by the outcome of the following optimization problem:

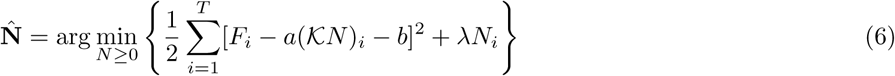

or equivalently:

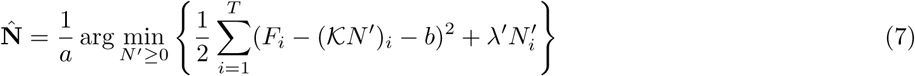

where λ, 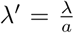 are *L*_1_ penalty coefficients that control the sparsity of the optimum (the higher *λ*, the sparser the optimum). When *λ* = 0 and the *N ≥* 0 constraint is relaxed, the optimal value 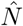 is exactly given by Eqn. 4. As shown in the next section, the choice of *λ* is crucial for efficient denoising and proper spike inference. Notice that the optimization problem is convex and can be solved efficiently in *𝒪* (*T*) for double-exponential kernels using the interior-point method, see [5]. This unusual linear scaling for a matrix inversion-like operation owes to the fact that *K*^*−*1^ is tridiagonal for double-exponential kernels: 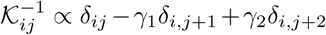, with 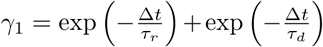 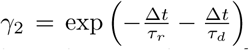, In [14], the authors apply the Pool-Adjascent Violator Algorithm originally developed for isotonic regression problems to solve this optimization in an even faster yet greedy way.

## II. DETERMINATION OF THE SPARSITY PRIOR *λ*

The choice of the regularization parameter *λ* is crucial. If it is too large, the inferred spike train is 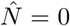 and all spikes are missed, whereas if it is too small, noise-induced transients are interpreted as spikes 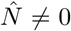, yielding large false positive rates. Intuitively, we expect the optimal choice to depend on the parameters of the generative model (noise level, spike amplitude, etc.) Here we review the expressions of *λ* previously used and we then introduce our method. We adopt the convention from Eqn. 7 and drop the primes. We assume for now that all generative model parameters are known.

### A Review of existing methods: fast-oopsi and constrained-oopsi

In [5], the authors derive the non-negative sparse deconvolution from an approximate Maximum A Posteriori principle. They assume that the spike count *N*_*i*_ at time step *i* follows a Poisson distribution of mean *v*Δ*t*, where *v* is the firing rate. After approximating the Poisson prior with an exponential distribution, they compute the negative log-likelihood *−* log *P* (*F, N*), which they find to be proportional to (7) with a sparsity prior *λ* given by:

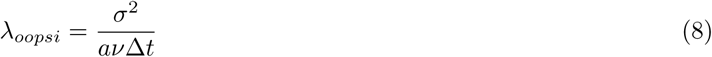

This approach thus provides an analytical expression for *λ*. However, due to the exponential approximation, this expression proves to be ineffective in several realistic experimental conditions as shown in Section 2C,D and illustrated in Figure 1.

To address this issue, a non-analytical method called constrained-oopsi (refered to as con-oopsi in the following) was recently introduced in [13]. The authors propose the following constrained deconvolution:

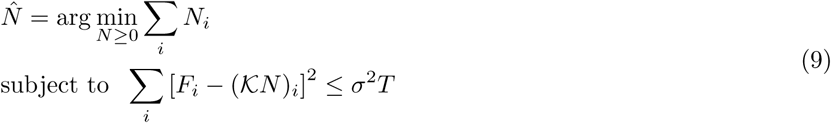

Where *T* is the number of observations. The problem can be rewritten using the Karush-Kuhn Tucker conditions by introducing the Lagrangian 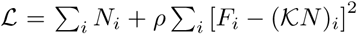 where *ρ* is the Lagrange multiplier associated with the constraint. There exists *ρ* such that the critical point *N*^*\**^ of *L* is the solution of the constrained optimization problem. Clearly, *N*^*\**^ satisfies the constraint only if *ρ* is non-negative; in this case *L* is convex and the critical point is a minimum of *L*. Overall, the optimization problem can be rewritten as:

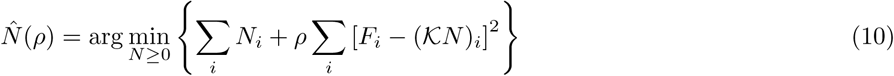

Identifying 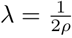, the constrained deconvolution is equivalent to a sparse deconvolution with an adaptive sparsity prior λ. Since 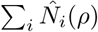 is a decreasing function of *ρ*, the expression for *λ* reads:

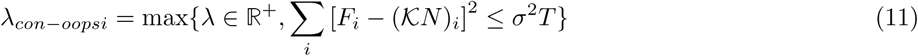

In practice, *λ*_*con−oopsi*_ is found by alternatively solving Eqn. 7 and updating *λ*, by decreasing it if the reconstruction error is too large, or increasing it otherwise. This non-analytical approach performs better than fast-oopsi, see Section 2C,D and Figure 1. However, it comes at a cost of increased computational time, because the deconvolution problem must be solved many times and the number of iterations required for convergence is not known in advance.

### B. Blind Sparse Deconvolution

We propose a different analytical expression for *λ*, inspired by [21]. It is deduced from the analysis of the optimization problem for two simple configurations, in which there is either zero or one spike in the original signal. We show that this solution combines the computational speed of fast-oopsi and the robustness of con-oopsi.

#### 1. Spikeless Signal

In the following, we use matrix notations and rewrite the cost function as:

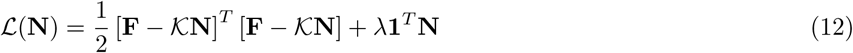

The gradient writes:

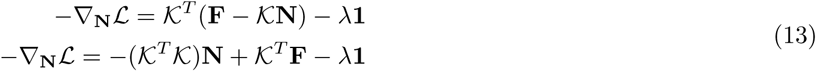

Let’s first assume that the signal is spikeless, such that *F*_*i*_ = *σ∊*_*i*_, where *∊*_*i*_ is a gaussian white noise. Since (*K*^*T*^ *K*)*N >* 0, we have:

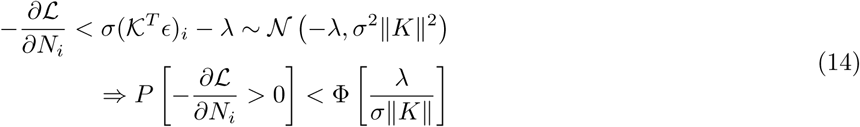

where 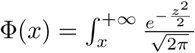, and 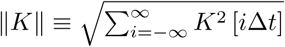

Therefore, if *λ* = *λ*_1_ *= z*_1_*σ∥K∥* with *z*_1_ large enough, the gradients are almost always negative, and the global optimum of *L* is 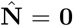 = 0. Hence for instance, setting z_1_ = 2.326, yields a probability of false positive event per time bin P_*F* *P*_ *<* 0.01.

#### 2. Single spike signal

We now examine a configuration in which a single spike is present in the data:

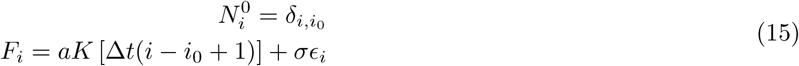

The gradient writes:

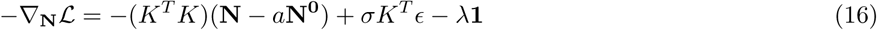

We look for an optimum of the form 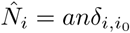 The optimization with respect to n gives:

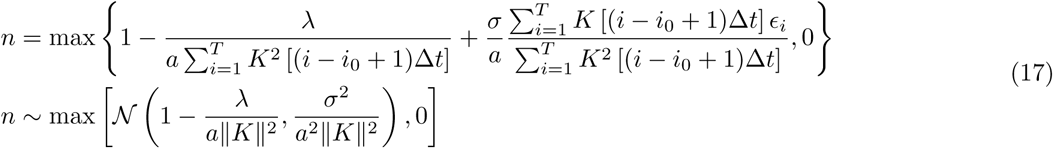

where the last line assumes that 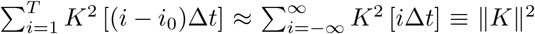, which is true provided that *i*_0_ is far from the boundaries. Thus, if the spike position is known in advance, the inferred spike is a thresholded gaussian variable.

One may observe that the noise-to-signal level 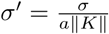 that appears in this expression is smaller than 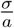 a factor 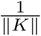. This has an important consequence: since max (*K*) = 1, the norm 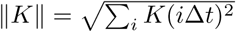 of the discretized kernel is proportional to 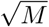 where *M* is the typical number of time frames over which *K* is non-zero. Thus, 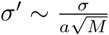 as if the noise had been averaged over the duration of the transient. This suggests that low SNR signals can be efficiently inferred provided that the spike-induced fluorescence transient is sufficiently well sampled.

Eqn. 17 shows that when *λ* is too large, the probability that a given spike is undetected reads:

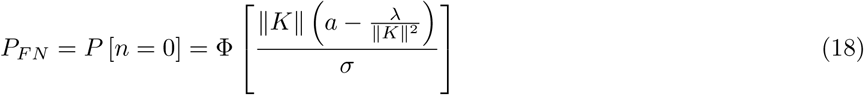

Therefore, setting 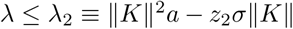, with, say, *z*_2_ = 2.326, guarantees a low false negative rate (FNR) as the probability that a spike is detected is then larger than 0.99.

The sparsity prior *λ*_*BSD*_ is chosen to minimize both the FPR and FNR. Hence, for *a* = 1, *σ* = 0.1, *τ*_*r*_ = 0.1, *τ*_*d*_ = 0.5, *f* = 10*Hz, λ*_2_ = 4.1379 is much higher than *λ*_1_. In this case, setting *λ*_*BSD*_ = *λ*_1_ is the best solution, as smaller values of *λ*_*BSD*_ lead to less signal deformation. In contrast, for configurations such that *λ*_1_ *> λ*_2_, *i.e.* when 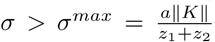, it is impossible to satisfy both constraints (low FPR and low FNR); in this case we use the crossover value 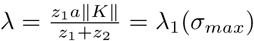.

#### 3. Sparsity prior for BSD

To summarize, in our Blind Sparse Deconvolution (BSD) algorithm, the sparsity prior is set analytically as:

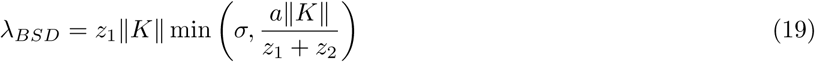

where 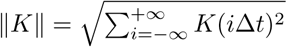 is the *L*_2_ norm of the discretized convolution kernel *K*, and *z*_1_, *z*_2_ are two numbers *∼* 2 that control the precision and recall, respectively.

#### 4. Thresholding the BSD inferred signal

Some applications, such as network connectivity inference, may require to threshold the signal in order to get a binary spike train. Unlike previous methods, BSD provides rationale for choosing a threshold. Indeed, the computations performed in Section 2B1,2B2 shows that the (unnormalized) inferred spikes in the absence (resp. presence) of spikes are thresholded gaussians variables, with means 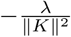 and 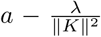, respectively, and identical variance 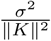 Picking a threshold that separates the two distributions yields:

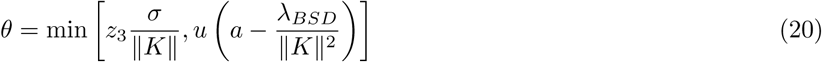

where *z*_3_ is a quantile of the normal distribution, and *u* a number between 0 and 1, say, 0.5. When *θ* equals the left term, the vast majority of the noise is efficiently filtered out such that any non-zero value in the output signal can be safely assigned to a spike; the right-hand term in turn prevents the threshold from becoming larger than the signal itself.

### C. Qualitative Comparison

As we have just shown, the sparsity prior expression *λ*_*BSD*_ allows one to simultaneously minimize both the FPR and FNR. In contrast, the expression of *λ*_*oopsi*_ offers no guarantee that either is small in all situations. For instance, if *a* = 1, *σ* = 0.1, *τ*_*r*_ = 0.1, *τ*_*d*_ = 0.5, Δ*t* = 0.1*s, v* = 1*Hz*, we find *λ*_*oopsi*_ = 0.1 and *λ*_1_ = 0.51; hence *λ*_*oopsi*_ is too small which results in multiple noise-induced false spikes. On the other hand, for *σ* = 0.25 and *v* = 0.1*Hz*, we have *λ*_1_ = 1.25, *λ*_2_ = 3.39 *λ*_*oopsi*_ = 6.25; in this case, *λ*_*oopsi*_ is too large, such that most of the spikes are missed.

We expect good performance as well for *λ*_*con−oopsi*_, although differences with *λ*_*BSD*_ may exist. In the absence of spikes, the noise is expected to be completely filtered out, because the optimum of Eqn. 9 is 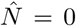 in the large *T* limit. In the presence of spikes, we expect the spikes train to be recovered to some extent, since 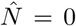 violates the constraint of Eqn. 9 but in a slightly noisier fashion. Indeed, observe that as soon as *λ >* 0, the spikes are on average underestimated: 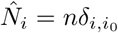 with *n <* 1, see Eqn 17. Thus, if there is a ‘good’ value *λ* such that we retrieve exactly the spikes at their positions, we typically find at the optimum 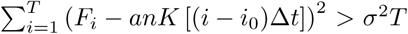. Hence the constraint is violated, and *λ* is decreased, until false (noise-induced) spikes appear and reduce the reconstruction error below *σ*^2^*T*.

As an illustration, we compare the results of the three inference algorithms for a signal with *σ* = 0.4, *f* = 10*Hz* in Figure 1a. For fast-oopsi, the sparsity prior is too large, and no spikes are inferred, whereas for both con-oopsi and BSD, the signal is correctly recovered. We notice however that BSD infers slightly less false spikes than con-oopsi.

### D. Quantitative Comparison

The performance of the three algorithms is now compared on a systematic benchmark. A random spike train is drawn from a Poisson distribution of mean firing rate *v* = 0.1*Hz* over a duration *t* = 10000*s*. The signal is generated with the discrete model Eqn. 3 and a noise *σ* of varying amplitude. We use a double exponential kernel *K* with *τ*_*r*_ = 0.1, *τ*_*d*_ = 0.5, akin to a GCaMP6 reporter. Spike trains 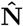 are inferred *with knowledge of the generative model parameters* and compared with the original spike train **N**.

#### 1. Precision-Recall

The precision-recall performance is evaluated as follows:

- The continuous spike train is binarized with a threshold *θ*.
- A spike is considered to be detected if the closest detected spike is within a time interval *δt* around the original spike; a true positive rate is computed (between 0 and 1) as the number of false negatives over the number of actual spikes (see [9]).
- Conversely, a false positive rate (FPR) is computed as the ratio of the number of inferred spikes that do not correspond to a real spike, over the number of actual spikes.
- The threshold is varied from *∞* to *θ*_*min*_, such that *θ*_*min*_ corresponds to a maximum acceptable FPR, here taken to 1.
- Precision-recall graphs are plotted, and the area under the curve is computed.

The results are shown in Figure 1b. We observe that the fast-oopsi algorithm performs sometimes very well (*f* = 100*Hz*, high SNR) sometimes equivalently as con-oopsi and BSD (*f* = 100*Hz*, lower SNR), but often very poorly (*f* = 10*Hz*); Such unreliability may be highly detrimental in actual experiments. BSD and con-oopsi compare equivalently, with BSD slightly outperforming con-oopsi in most configurations. Notice that at low sampling rate (*f* = 1*Hz*), the P-R curve saturates at higher levels for BSD, suggesting that a large fraction of the spikes go undetected with con-oopsi.

#### 2. Speed

BSD and fast-oopsi share the same algorithms, albeit with different parameters values; hence they have similar computational cost. The con-oopsi implementation, on the other hand, is slower because the sparse deconvolution has to be performed many times with different values of *λ* until convergence is reached. In practice, for experiments performed on a MacBook Air 2013, with 1.3 GHz Intel Core i5, we find a 3 to 25-fold increase in computation speed, depending on the array size. Our experiments shows that the number of iterations can be suprisingly large in practice. In particular, if the noise level is underestimated by con-oopsi, the error constraint is tighter and adding the positivity constraint may lead to no solutions at all - yielding many iterations in vain and increased computational time, see Table I. This reflects in the fact that the computing time is largely dependent on whether or not the noise level is provided.

Notice that the exact gain in speed depends on which version of con-oopsi is used (here, Matlab implementation, con-oopsi version of Dec. 2015, with cvx). Although we did not test the PAVA optimizer, we expect a gain of the same order of magnitude between constrained-PAVA and BSD-like PAVA. Such a difference in computation load may prove highly beneficial for real-time inference in high data-throughput recordings, as illustrated in section 6.

## III. THEORETICAL LIMITS ON THE PRECISION-RECALL AND TEMPORAL RESOLUTION

The fundamental motivations for spike train inference are to denoise the fluorescence signal and to improve the temporal resolution of the neural recording. Theoretically, if the generative model is correct, the convolution kernel is known and the isgnal is noiseless, then perfect retrieval of the spike train in terms of detection and timing can be achieved. Because of the noise, the accuracy is in practice limited by the rise and decay times: some spikes can be missed, have a wrong timing or be split across two successive time bins. These limitations have been characterized quantitatively in [22] in the context of Bayesian inference, when the noise is Poisson-like, *τ*_*r*_ is negligible and without super-resolution. However, no such analysis has been performed for sparse deconvolution algorithms. The BSD package incorporate routines to compute theoretical true and false positive rates and temporal resolution, for any given set of parameters (*a, σ*, **K**) that can be extracted from the data.

### A. Precision-Recall for isolated spikes

The theoretical false positive and negative rates (FPR, FNR) are first computed within the sparse deconvolution framework with *λ*_*BSD*_. For the false positive rate, the computation was performed in Section 2B: we obtain a probability of false positive rate per time bin:

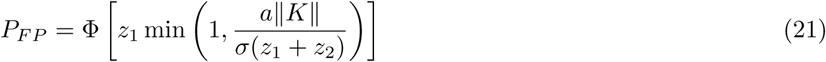

For the false negative rate, we follow a similar reasoning than in Section 2C: we consider a signal of the form *F*_*i*_ = *an*_0_*K* [Δ*ti − t*_0_]+ *σE*_*i*_ with *t*_0_ = (*i*_0_ *−* 1)Δ*t* + *δt*_0_ and 0 *≤ δt*_0_ *<* Δ*t*. Note that we have now relaxed the previously made approximation *K* [Δ*ti −t*_0_] *≈ K* [Δ*t*(*I − i*_0_ + 1)] in order to probe the effect of intermittent sampling. We obtain a lower bound [32] for the probability of false negative per spike:

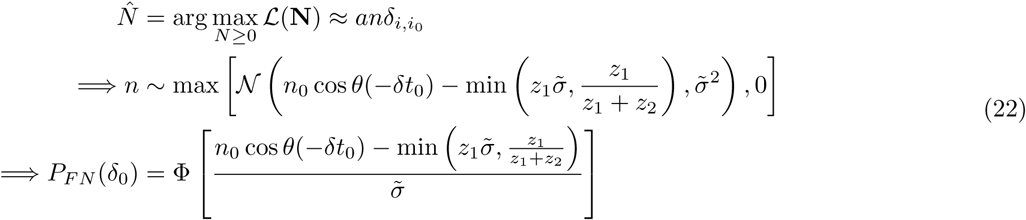

Where

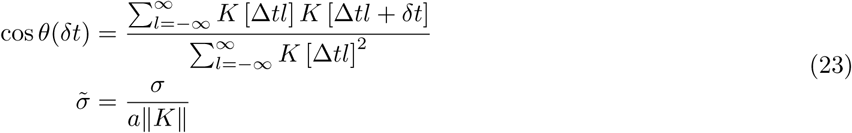

Note that the probability depends on *δt*_0_; for instance if *τ*_*r*_ = 0 and *δt*_0_ *<<* Δ*t*, spikes emitted right after a measurement yield low-amplitude fluorescent transients and are thus likely to be missed. Overall, the probability of false positive is given by:

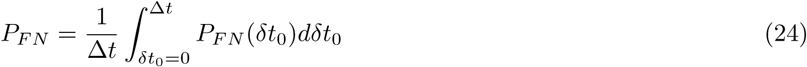

We display in Figure 2(a) the true positive rate (TPR) for different sampling rates, as a function of the noise level for *τ*_*r*_ = 0.1, *τ*_*d*_ = 0.5. The FPR is set at 0.01/frame (i.e. *λ*_*BSD*_ = *λ*_1_ and *z*_1_ = 2.366). An important insight of this graph is that at low sampling rate, the TPR quickly decays with the noise level because spikes emitted shortly after a measurement are often completely missed. Conversely, improving the TPR (with e.g. *z*_2_ = 2.366) yields a large number of false positives at low sampling rate.

**FIG. 2:**
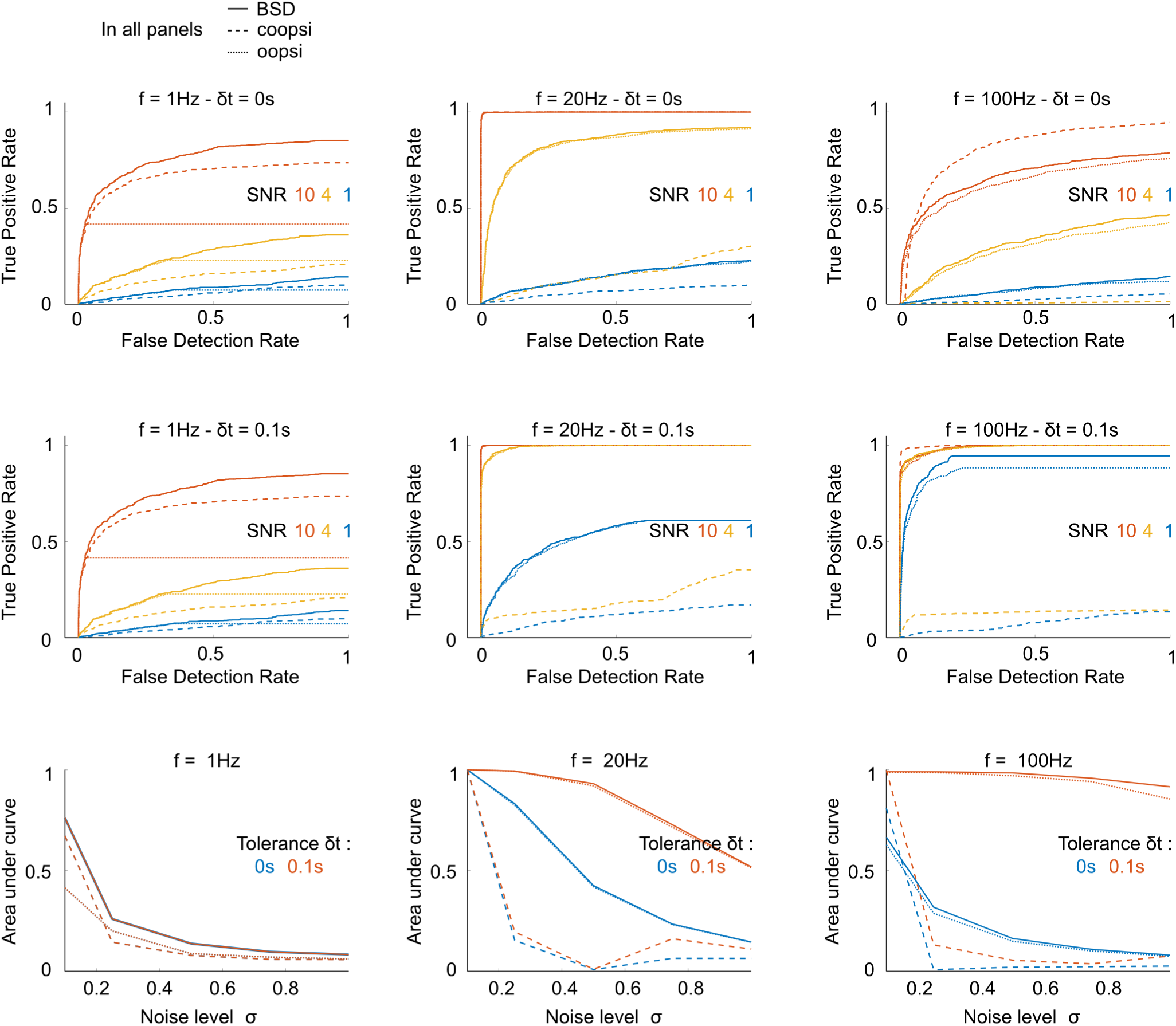
*f* = 10*Hz, σ* = 0.4, *τ*_*r*_ = 0.1, *τ*_*d*_ = 0.5. Precision-recall curves and area under the curve. Top row: PR curves for *f* = 1, 20, 100*Hz*, and SNR, with a time tolerance for spike detection of *δt* = 0. Middle row: same, with time tolerance *δt* = 0.1*s*. Bottom row: Corresponding area under the curve as function of the noise level.

### B. Temporal Resolution

#### 1. Rough analytical estimate for isolated spikes

We now estimate the temporal resolution by examining the probability distribution of 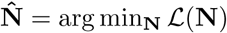 = arg min**_N_** *L* (**N**) given a single-spike noisy fluorescence signal *F*_*i*_ = *aK* [Δ*t*(*i − i*_0_ + 1)] + *σE*_*i*_ [33]. Since the distribution cannot be computed explicitly, we start with a simpler heuristic computation, and study the distribution of the initial negative gradients 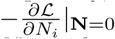 Consider indeed the gradient descent optimization dynamics. Because of the *L*_1_ penalty, large components *N*_*i*_ tend to grow faster and to screen neighbouring small components, yielding sparse solutions with only few non-zero components; it is therefore likely that the largest components of **N** after one gradient descent step (after which 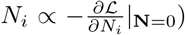) remains the largest at the end of the optimization. Hence if the initial negative gradient is larger at position *i*_0_ + *δ* than at position *i*_0_, we expect the inferred spike 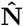 to be similarly delayed with respect to the true spike position. The probability of such an error can be computed as:

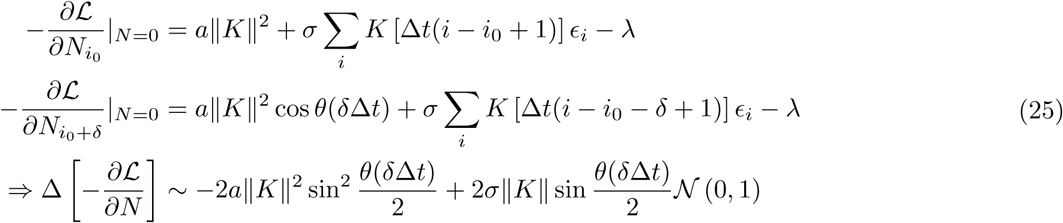

where the angle *θ*(*δt*) is defined in Eqn 23.

Thus, the initial gradient at the offset time *i*_0_ + *δ* is higher than its value at the spike time *i*_0_ with probability 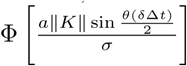, typically resulting in a time-shifted inferred spike. This results in a typical timing error *δt* on the spike position of the order of:

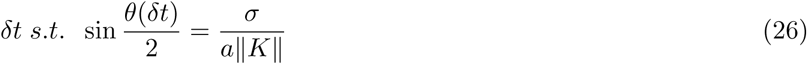

This timing error is a non-trivial function of the kernel *K* and the noise level. The higher the effective noise level 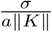, the higher *δt*. The second factor is small for rapidly growing *θ*(*δt*), *i.e.* when the overlap between the kernel *K*(*t*) and its lagged version *K*(*t* + *δt*) is a fast decaying function of *δt*. Hence, the ‘sharper’ the kernel, the lower *δt*. Notice that *δt* does not depend on *λ*, suggesting that the temporal resolution is intrinsically limited by the noise, and not by the algorithm.

#### 2. Response function for isolated spikes

In a more rigorous fashion, one can estimate the response function to an isolated spike under the assumption that the solution of the optimization is a Dirac, 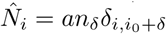 The optimization result is given by the following equation:

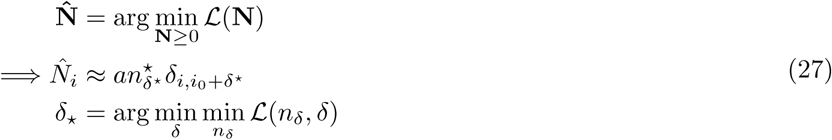

For a given fluorescence trace, the inner optimization over *n*_*δ*_ can be carried out analytically, and the outer optimization numerically. By computing the optimal offset *δ*^*\**^ for various noise realizations, one can estimate the probability distribution of the inferred spike offsets *P* (*δ*^*\**^). It is a function of the parameters *a, σ, K, z*_1_, *z*_2_. This computation can be easily generalized to support the super-resolution setting (see Annex C for the analytical details). Some response functions are displayed in Figure 2b-inset for typical calcium indicators. We used the parameters: (i) GCaMP6s: *τ*_*r*_ = 180*ms, τ*_*d*_ = 0.55, (i) GCaMP5k: *τ*_*r*_ = 58*ms, τ*_*d*_ = 0.52*s*, (i) GCaMP6f: *τ*_*r*_ = 25*ms, τ*_*d*_ = 0.38*s*, (i) OGB1-like: *τ*_*r*_ = 20*ms, τ*_*d*_ = 80*ms*. For the first three set of time constants, the values are deduced from the fluorescence recordings on mice V1 cells reported in [18].

We also display in Figure 2b the width *δt* (measured by a gaussian fit) as a function of the noise level, for a sampling frequency *f* = 60*Hz*. As expected, the temporal resolution of the spikes can be lower than the sampling period if the noise is large, and we observe that reporters with large *τ*_*r*_ yield lower temporal resolution.

We finally examine the impact of the sampling frequency on the temporal resolution. In an experiment with a fixed number of sampled neurons, increasing the sampling rate *f ≡* Δ*t*^*−*1^ by a factor s typically comes at the cost of reducing the exposure time *τ*_*e*_ by the same factor *s*, which in turn increases the noise *σ* by 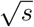 it is therefore not obvious that one would improve the temporal resolution by increasing the sampling frequency. We display in Figure 2c for the same set of calcium indicators, the inverse width *δt−*^1^ as a function of the sampling rate *f*, for various signal to noise ratios (SNRs) at a reference frequency 10 Hz. We see that *δt*^−1^ saturates at a value that depends on the SNR and on the rise and decay constant times. Hence, with GCaMP6s, increasing the frequency beyond 50 Hz does not result in improving the temporal resolution.

The fact that the temporal resolution saturates can be seen from Eqn 26: asymptotically, we have 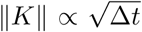, and since 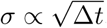, the effective noise level 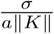 reaches a well-defined limit - and so does *δt*.

## IV. HYPERPARAMETERS LEARNING

All sparse devonvolution methods rely on the knowledge of the generative model’s parameters. However, owing to the variability in the calcium reporters intracellular concentration and other biochemical cellular processes, these paramaters may significantly vary from experiment to experiment, and for different neuronal types. In the fast-oopsi implementation, the authors proposed to infer the parameters (*a, b, σ, v*) in an iterative way: an initial guess is made, deconvolution is performed, parameters values are then updated based on the deconvolution result, whereas for con-oopsi, the authors propose to estimate *σ, K* only once. We follow the same iteration-based approach as fast-oopsi, but the parameters are inferred and refined differently; we also add a method to infer and refine the kernel *K*.

### A. Initial estimation of the parameters

We are given a time series of the form **F** = *a K* **N** + *σE* + *b*, with unknown *a,b,σ, K*. In the following, we assume that the baseline is constant or equivalently that the variable baseline has been previously estimated and subtracted from the signal. From the knowledge that *N* is non-negative and sparse, we deduce that:

- The baseline *b* is essentially the most often observed value of *F*; the data histogram is computed, and *b* is estimated as the center of the interval with highest frequency. Using the median of *S* also provides a good estimator.
- All activity below the baseline originates from the noise, hence *F* = *F* [*F < b*] *− b* follows a half-Gaussian distribution min [*N(*0, *σ*^2^),0];it is fitted to deduce *σ*.

In a similar spirit to con-oopsi, we estimate the convolution kernel *K* through the signal auto-correlation matrix. Indeed, observe that:

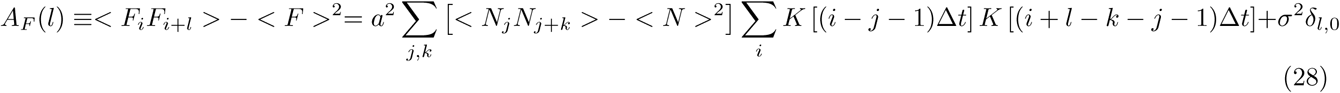

Under the assumption that the spiking events *N*_*i*_ are independent, identically distributed Poisson variables, we have 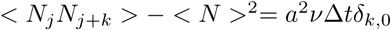 and Eqn. 28 can be simplified as:

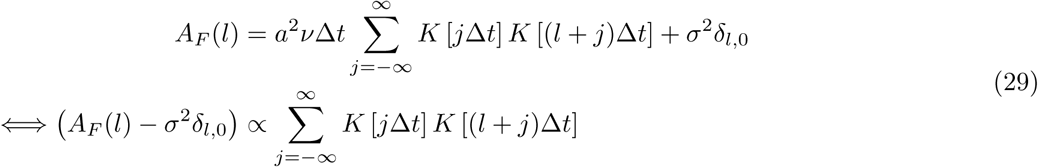

The auto-correlation matrix can be estimated from the data as 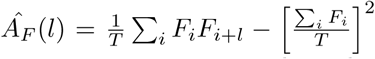. Together with the previous estimate of *σ*, the left-hand side of the equation can thus be estimated [34]. The right-hand side is the overlap between the kernel *K* and its delayed version *K*^*I*^(*t*) = *K*(*t* + *l*Δ*t*). We can normalize both terms to 1 for *l* = 0, and use a least square fit to estimate *K*.

Lastly, the spike amplitude *a* and frequency *v* can be deduced from the following equations, that hold under the model assumption:

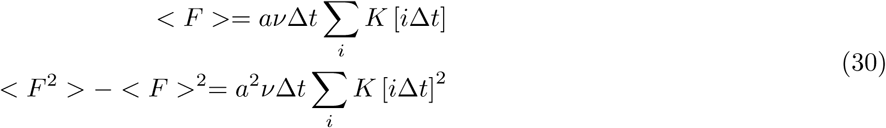

Although they yield very good results for synthetic datasets, these estimators can fail in several frequently encountered situations in practice:

- When the neural activity is not sparse, we do not expect *b* to be the most frequent fluorescence value. An error in the estimation of *b* can result in a misestimation of *σ* as well.
- When the neuron displays bursting activity (*i.e.* several spikes in short time intervals), the hypothesis that the *N*_*i*_ are independent usually fails. This may result in overestimating *τ*_*r*_ and/or *τ*_*d*_.
- In the same situation, Eqn 30 is incorrect and *a* can be overestimated.
- When the noise exhibits temporal correlation (streaking artefacts in light sheet imaging, small sample drifts, fluctuations in laser intensity, etc.), the white-noise hypothesis does not hold, which may result in a misestimation of *τ*_*r*_ and *τ*_*d*_.

We systematically studied the bias in spike inference that arises when the estimated time constants *τ*_*r*_ and *τ*_*d*_ differ from their true values, 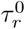 and 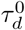. As illustrated in Figure 3a-d, inferring the spikes with an incorrect convolution kernel leads to systematic errors. Suppose for instance that 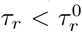 and 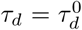 (Figure 3a). Then a spike-induced fluorescence transient tends to exhibit a faster initial rise than expected. Hence, from a Bayesian perspective, such a transient is likely to be interpreted as two small consecutive spikes. Such a mismatch in the kernel parameters will therefore show up as duplicate spikes. In general, the nature of the error depends on the kernel mismatch:

**FIG. 3:**
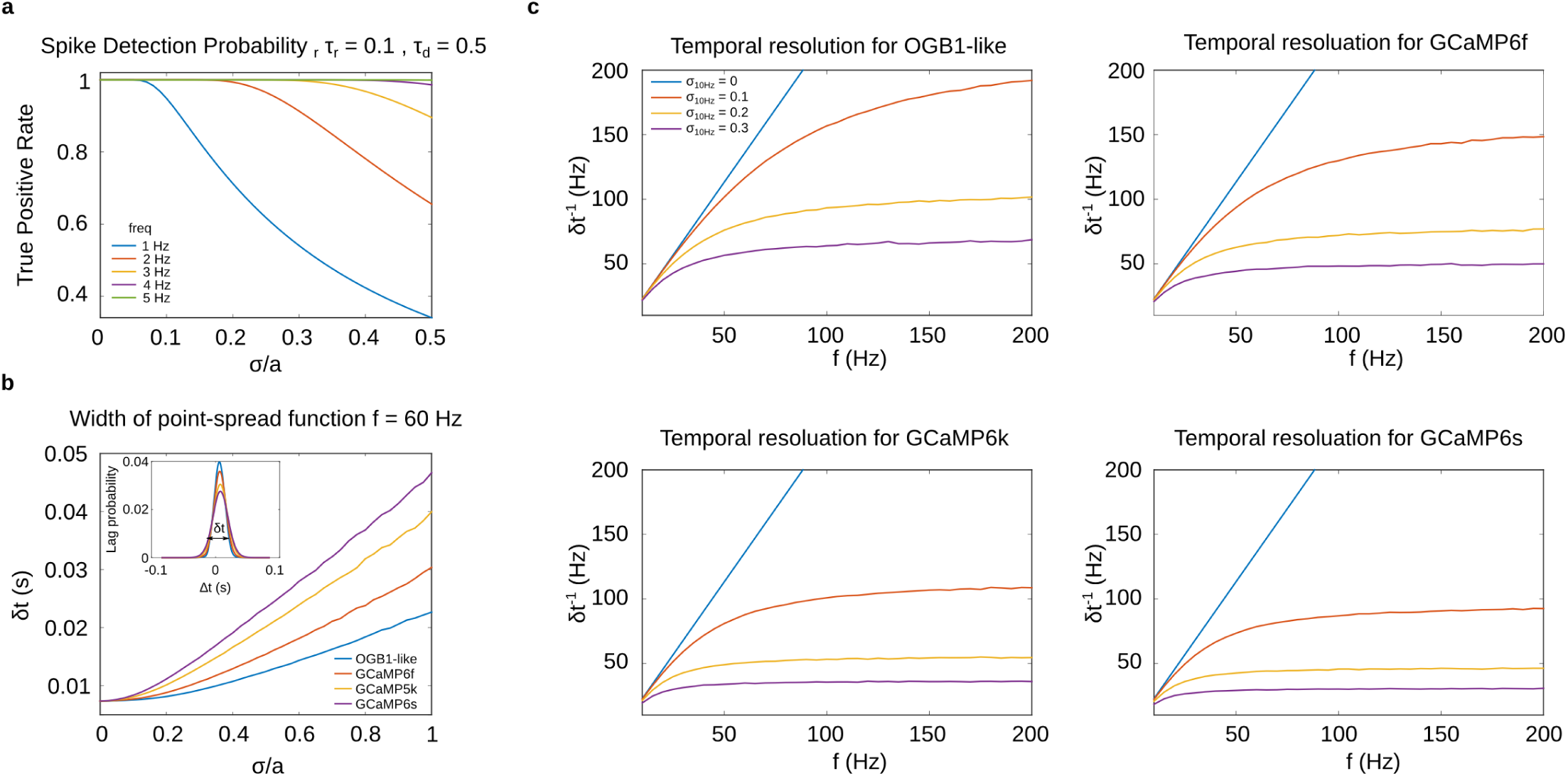
(a) True positive rate of BSD as a function of the noise level for different sampling frequencies, for a fixed FPR of 0.01/frame, (b) Width of the response function as function of noise for various calcium indicators, at fixed frequency *f* = 60*Hz*, (c) Width of the response function as a function of the sampling frequency for various calcium indicators and reference noises.

**FIG. 4:**
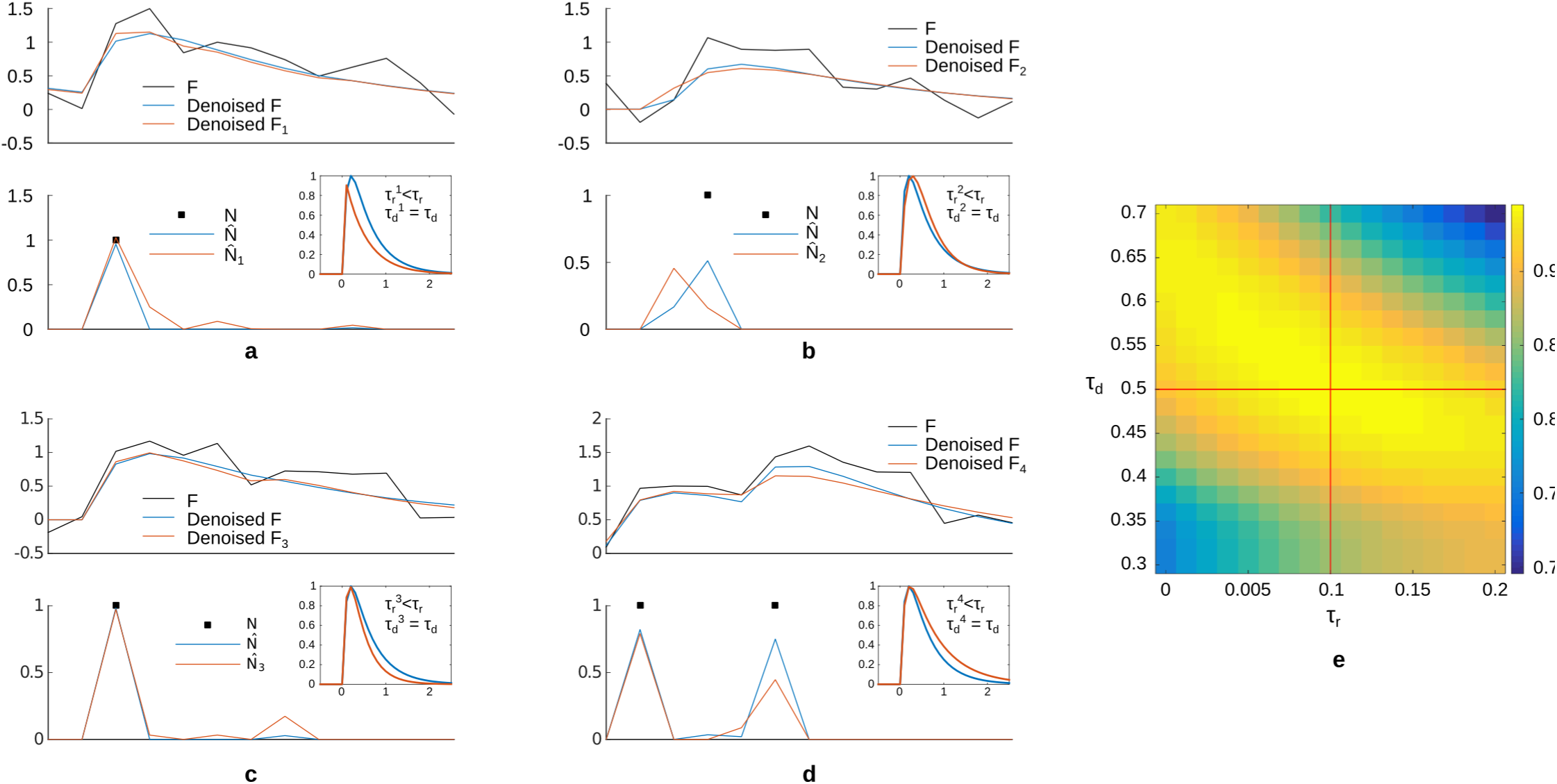
Left: Example results of spike inference on synthetic data with mismatched convolution kernels. For each of the four figures, a fluorescence signal is generated with a kernel *K*^0^; inference is performed with true parameters (blue curves) and with mismatched parameters (red curves). The true and mismatched kernels *K*^0^ and *K* are depicted (insets). Systematic errors appear in the spike timings. Right: Area Under Curve classification performance with time tolerance *δt* = 0*s*, as a function of the rise and decay time constants. The parameters used to generate the signal, depicted in red, are typical of a GCaMP6 reporter.

- 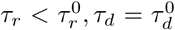 (Figure 3a): the inferred spikes are split in two, to compensate for the smaller rise time than expected for a single spike.
- 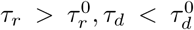 (Figure 3b): the inferred spikes are in advance, to compensate for the faster rise of the fluorescence signal.
- 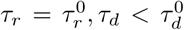 (Figure 3c): the inferred spikes exhibit ‘echos’ to compensate for the slower than expected decay of *F*.
- 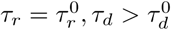 (Figure 3d): the inferred subsequent spikes are ‘screened’ (lower amplitude) to compensate for the slower than expected signal decay.

We quantified how a kernel misestimation degrades the decoding performance by evaluating the relative reduction in precision-recall (area under curve) for various offsets of *τ*_*r*_ and *τ*_*d*_ (Figure 3e). Interestingly, some direction of the mismatch vector can be less deleterous: when both 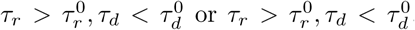, the loss in performance remains modest.

### B. Iterative parameter estimation: adaptive blind deconvolution

As previously explained, the initial generative model parameters are not always the true ones. To improve the estimates, let us write the cost function:

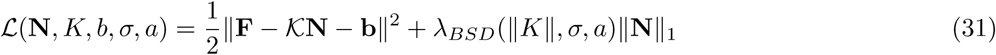

The spikes are inferred by optimizing *L* with respect to **N**. In order to improve the model parameters, we also optimize over *K* and *b* through a coordinate gradient descent:

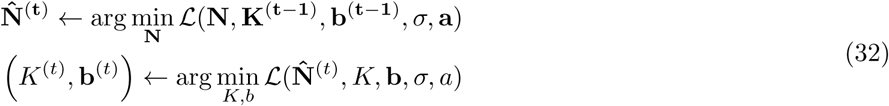

The first optimization step was discussed in Section 1. The second optimization is a parametric temporal regression problem; it can be solved efficiently in 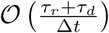 by introducing the cross-correlation 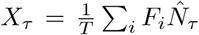 and auto-correlation 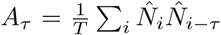 functions up to some cut-off 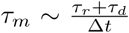 (see details in Annex B). The point of this step is that if *N* ^(*t*)^ = *N*_0_, then the optimum is exactly *K*_0_ if *σ* is small or *T* is large. More generally (*N*_0_, *K*_0_) is a fixed point of the optimization dynamic in the low noise limit, and intuitively, we expect that at finite noise, another fixed point close to (*N*_0_, *K*_0_) exists and can be reached. We show in Annex B that *K*^0^ is the global optimum in the case of isolated spikes and low noise level. The optimization will not necessarily converge to such solution because the function *L* (*N, K*) is not convex, and only local minima are found. In practice, the optimum is usually very close to the original convolution kernel, and is reached if the initial estimate is good enough (Figure 5). The convergence can be improved by thresholding the spikes before updating the kernel, as it prevents false spikes from contributing to the cross-correlation. The iterative process is no longer an optimization but it still converges.

**FIG. 5:**
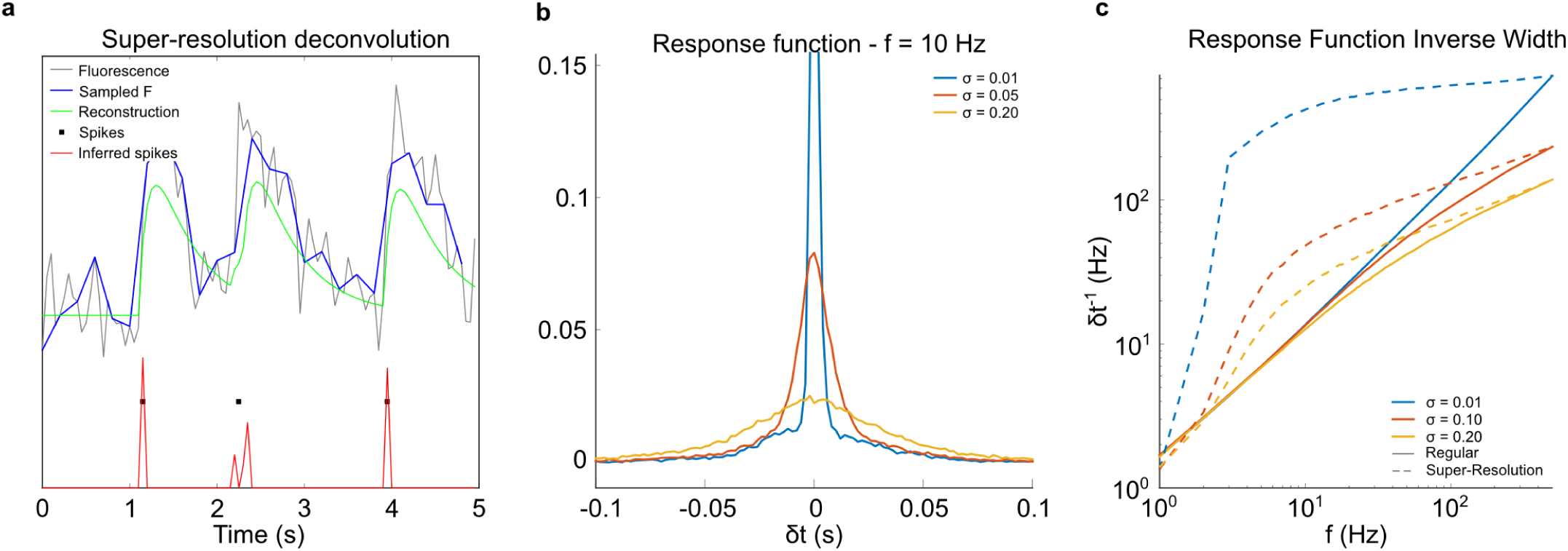
(a) Example of super-resolution inference: a fluorescence signal is generated at *f*_0_ = 20 Hz, sampled at 5 Hz and reconstructed at 20 Hz. Parameters: *τ*_*r*_ = 0.1, *τ*_*d*_ = 0.5, *σ* = 0.2, *a* = 1. (b) Response function at 10Hz for various noises. Notice a smaller width than the sampling interval 0.1s (c) Inverse width *δt*^*−*1^ of the point-spread function, as function of the sampling frequency for regular reconstruction (full) and SR reconstruction (dotted).

The noise *σ* and spike amplitude *a* can be refined as well, using

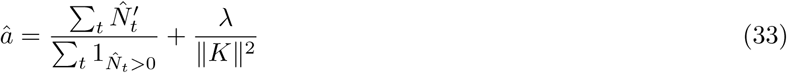

Where the last term corrects the bias due to the sparse prior (see Eqn. 17)

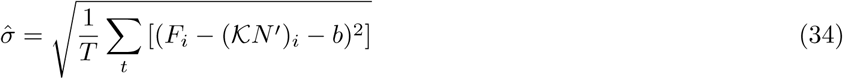

## V. BEYOND THE DISCRETE MODEL: TEMPORAL SUPER-RESOLUTION

Most fluorescent microscopy techniques –two-photon, confocal or scanning light-sheet– involves the sequential scanning of laser beam at different locations within the sample. Hence, for a given dwell time of the laser at each neuron position, there is a trade-off between the sampling rate and the total number of sampled neurons. In other experimental fields, resolution limitations due to recording constraints have been significantly circumvented through signal processing algorithms. For instance, super-resolution microscopy achieves imaging at higher resolution than the diffraction limit [23–27], and compressed sensing applied to MRI allows to drastically reduce the number of measurements required to reach a given resolution [28]. These algorithms rely on the hypothesis that the original signal is sparse in a certain basis - it is therefore tempting to apply them to our problem, given that neural spikes are sparse in the canonical basis. This possibility had been discussed in the context of bayesian inference [4]. Temporal resolution was shown to be slightly improved in very specific settings, *i.e.* when using prior knowledge of inputs (stimulus) and spiking history dependence of the neuronal activity. In this section, we demonstrate on synthetic data how the blind sparse deconvolution framework can be extended to support super-resolution in more generic contexts.

### A. Qualitative analysis

We start off with a qualitative analysis and consider the fluorescence signal produced by an isolated single spike of amplitude *n*:

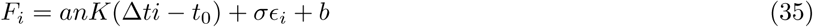

Denoting 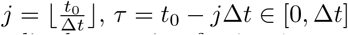, 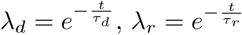 and assuming for simplicity that *b* = 0 and that *K* is unnormalized, we write, for *i > j*:

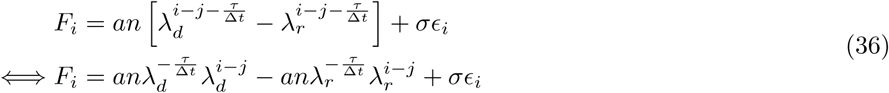

Thus, the observed fluorescence is a double exponential with non-equal coefficients of the form 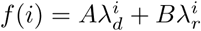 Fitting the coefficients with a least-square method yields estimates of 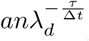 and 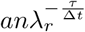, which can be converted to estimates of *τ* and *n*. Thus, it is possible in principle to find the exact spike position in the noiseless case, if we know a priori that the signal contains a single spike. Notice that this is possible only if *λ*_*r*_ *>* 0, *i.e. τ*_*r*_ *>* 0; if *τ*_*r*_ = 0, the observed fluorescence is a single exponential of amplitude 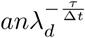, and we cannot recover both *n* and *τ* without ambiguity [35]. In the case of a noisy signal, we expect that super-resolution can be achieved only if 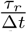 is large enough with respect to some function of *σ*. Notice also that if multiple spikes occur within the same time bin, the observed fluorescence transient is still a double exponential with non-equal coefficients, and it cannot be distinguished from the one produced by a single large spike at some average position. More generally, resolving two spikes in the same time bin would require the use of more complex convolution kernels.

### B. Generative model

With these limitations in mind, we now extend the deconvolution framework to implement super-resolution. The fluorescence signal is constructed using a discrete generative model at a fine-grained time scale 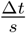, where *s* is a non-zero integer, which is then down-sampled by the same factor *s*. This yields the following generative model:

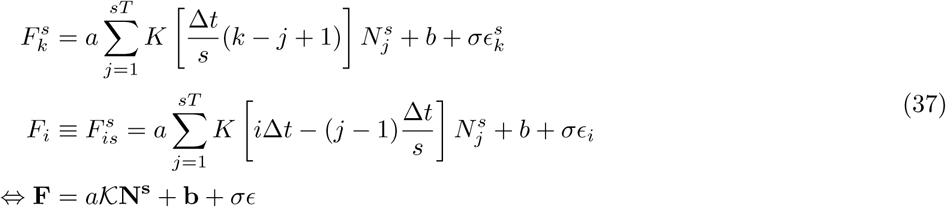

where *F*_*i*_ is the fluorescence measurement at *t*_i_ = *i*Δ*t* and *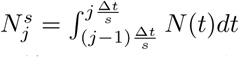 N* (*t*)*dt* is the number of spikes emitted in the time interval 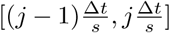. [36] The convolution matrix *K* is now rectangular, of size *T × sT*. It is not translation invariant anymore with respect to the spikes index *j* as the norm of the transient, 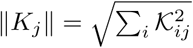 now depends on *j*. Indeed, writing *j* = (*p −* 1)*s* + *r*, we have:

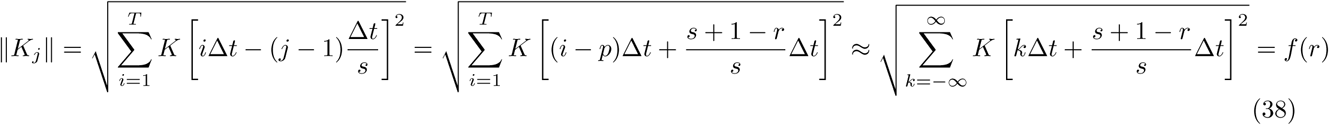

Typically, spikes occurring right after a fluorescence measurement (small *r*) have smaller *K*_*j*_ than spikes occurring right before a measurement (large *r*).

### C. Sparse Deconvolution

A sparse deconvolution algorithm is applied to estimate the spikes *N*^*s*^:

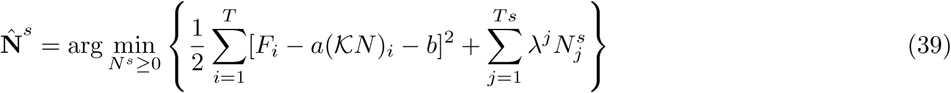

Notice that, although *K* is not invertible anymore, the optimum is still well-defined because of the sparsity penalty and non-negativity constraint. Compared to Eqn. 6, the main difference is that *λ* is not uniform anymore: 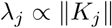 This property has an important consequence, as can be seen by considering the limit case *τ*_*r*_ = 0, *σ << a*. As discussed previously, a transient observed for *I ≥ i*_0_ can be interpreted either as a small spike right before the *i*_0_ measurement, or a ‘large’ one right after the *i*_0_ *−* 1 measurement. Thus, using a constant *λ* would systematically select the small spike interpretation, *i.e.* the inferred spike train would be systematically delayed with respect to the original spike train. This behavior is not desirable, and we would rather have both solutions to be degenerate global optima. This can be achieved by setting *λ* to a smaller value right after the *i*_0_ − 1 measurement. We show in Annex C that both efficient noise filtering and unbiased estimation of spike timing for isolated spikes can be obtained with the following expression for 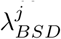

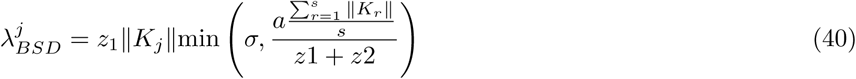

In practice, the optimization can also be performed efficiently using the interior-point method [5]. Adding a small *L*_2_ penalty 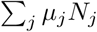, with 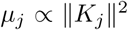 often provides better conditioning of the hessian, and faster convergence. It also ensures the unicity of the solution, in particular when *τ*_*r*_ = 0. Figure 5a shows an example of reconstruction of a signal generated at *f*_0_ = 20*Hz*, and sampled at 5*Hz*. We observe a good agreement with the original spike train; we observe in particular that, in spite of the sparse sampling, the onsets of the green and dark curves transients are very close to one another.

### D. Numerical experiments

We now test our algorithm on synthetic data generated using the model 37 at *f*_0_ = 500*Hz*, with *τ*_*r*_ = 0.1, *τ*_*d*_ = 0.5, spike frequency *v* = 2*Hz*. The fluorescence signal is down sampled to recording frequencies ranging from *f* = 1*Hz* to 500*Hz*. Spike trains are inferred with and without super-resolution. For super-resolution, we use 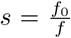, in order to reconstruct a spike train at the original frequency *f*_0_. For the case without super-resolution, an inferred spike train 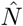 is first obtained at the sampling frequency *f*; to compare it with the original spike train at *f*_0_, we construct a signal 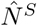 at sampling frequency *f*_0_ inferred signal by splitting evenly the spike counts:

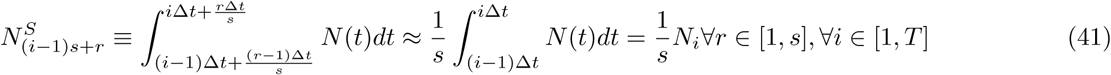

Once the signal is inferred, we measure the performance of the reconstruction in terms of temporal accuracy. A simple measure would be the cross-correlation between the spikes and the inferred spikes:

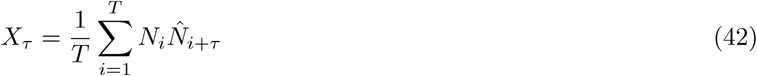

The faster *X*_*τ*_ decays to 0 as *| τ |* increases, the more accurate the reconstruction. This simple measure works well for independent spikes, but it does not produce the expected results when the spikes are temporally correlated: in the best case scenario, the spike is perfectly recovered 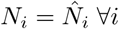, and we would have 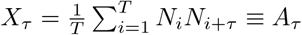. Since real spike trains may have significant temporal correlations, we introduce an estimator suited both for synthetic and real data sets, whose output does not depend on the spikes correlation. We define a response function *R*_*τ*_ through the temporal regression model:

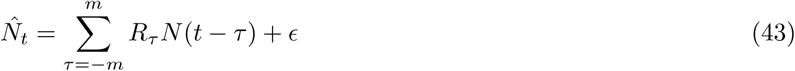

It can be estimated as:

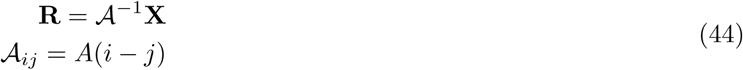

where *R*_*τ*_, *X*_*τ*_ are indexed formally as vectors **R**, **X**. *R*_*τ*_ is estimated for various sampling frequencies and noise levels, and results are depicted in Figure 5 b,c. They demonstrate that super-resolution is perfectly workable at small noise levels, and that significant resolution gain can be achieved at intermediate noise level typical of actual experimental conditions. For instance, at *f* = 10*Hz, SNR* = 5, the response function width is *∼* 2 *×* smaller than without super-resolution. Figure 5c shows that significant gain in resolution can be achieved as soon as the *f* 4*Hz*. This behavior is expected, as no gain can be achieved when 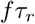 is small, see Section 5 A.

## VI. EXPERIMENTS ON REAL DATA SETS

We now apply our Blind Sparse Deconvolution algorithm to real data, and compare its performance with con-oopsi.

### A. Joint Electrophysiology and Fluorescence recordings on mice data

We first test it on joint electrophysiology and fluorescence recordings, as the former can be used as a ground truth for comparison. Recordings were performed on mice visual cortex at the Svoboda laboratory [17, 18], using the GCaMP5k, GCaMP6s and GCaMP6f probes, see summary in Table II. [37].For all datasets, the fluorescence is recorded at 60*Hz*, and the electrophysiology at 10*kHz*. For each neuron, the fluorescence signal is computed by simple averaging of the raw voxel-scale fluorescence over a region of interest. A baseline is computed using a moving percentile (window: 10*s*, quantile *q* = 0.15); it is subtracted to the fluorescence trace to remove the temporal fluctuations of the baseline. The resulting pre-processed signal is then fed to the following deconvolution algorithms:

- con-oopsi.
- BSD without iterative parameter estimation.
- adaptive BSD (up to 200 iterations). Each neuron is endowed with its own rise and decay times constants.
- BSD with super-resolution and adaptive parameters (up to 200 iterations). We use *s* = 5, i.e. spikes are reconstructed at 300 Hz.
- con-oopsi, BSD, and super-resolution BSD with ‘ground-truth’ parameters learnt using the spike positions, see below.

For each fluorescence probe, we investigate the spike detection precision-recall and temporal accuracy. For the former, we use the area under curve metric defined in Section 2 (Time tolerance: 1*/*60*s*); the AUC is computed separately for each recording, and a weighted average is calculated, with weights equal to the number of spikes per recording. For the latter, we estimate for each recording the response-function and compute the average; an offset and a width are estimated through a gaussian fit. The results are summarized in Tables III, IV and Figure 6.

**FIG. 6:**
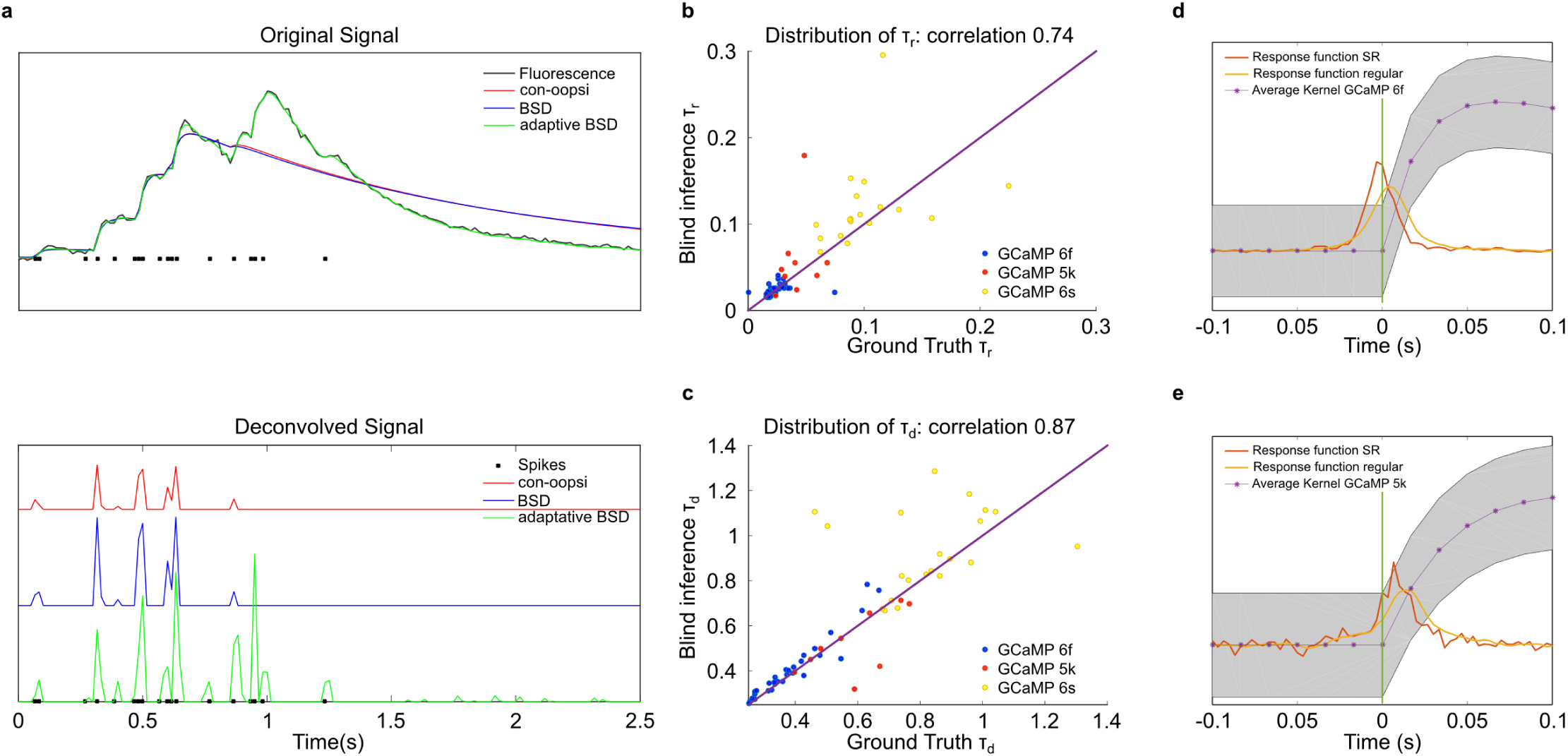
(a) Results of con-oopsi, BSD and adaptive BSD on joint GCaMP6f fluorescence and EP recordings of mice [17, 18]. (b,c) Distribution of inferred rise and decay time constants (d,e) Average response function for GCaMP6f/GCaMP5k recordings as function of the time to the spike onset. The inference is performed with or without super-resolution. The discrete kernel with median rise time, decay time and signal to noise ratio is displayed for comparison.

The inference is also performed with ‘ground truth’ parameters *τ*_*r*_, *τ*_*d*_, *b* for each neuron. The latter are obtained by minimizing the square error, using the knowledge of the position of the spikes provided by electrophysiology, (discretized at 60Hz): [38]

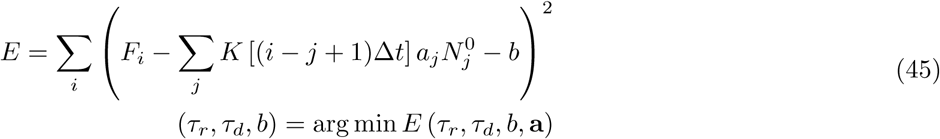

As expected from section 3A, the performance of the inference strongly depend on the inferred kernel. For the precision-recall, the main finding is that adaptive BSD always outperforms non-adaptive BSD and con-oopsi. Indeed, the uncorrelated spikes hypothesis used to estimate the kernel can be very wrong in practice, as illustrated in Figure 3a. Using the initial kernel estimates, con-oopsi and BSD misinterpret the fluorescence burst as being evoked by just a few spikes and a slow convolution kernel, whereas adaptive BSD correctly recovers the fast kernel and thus the true spike train. For all indicators, we found that inferred parameters are very similar to their ground truth values *τ*_*r*_, *τ*_*d*_ (figure 3b), such that using the latter for the inference yields little to no improvement in performance. When using the same kernel parameters, con-oopsi and BSD in general display comparable performance. In the absence of adaptive kernel estimation, the sparsity prior *λ*_*BSD*_, which depends on *K*, can be misestimated, resulting in weaker performance than con-oopsi. This advocates for the use of adaptive parameter estimation for robust inference.

In terms of temporal resolution, we find that GCamP6f reporter yields higher accuracy than GCaMP5k and GCaMP6s owings to its smaller rise and decay time. The measured temporal resolution can be compared quantitatively against the theoretical width depicted in Figure 2b, using the average measured noise to signal levels. The overall agreement is good, but the experimental width are *∼* 30% larger than the theoretical bounds. This is likely due to the fact that real spikes are not isolated, and neighboring spikes tend to decrease the temporal accuracy.

We find that the super-resolution improves the temporal resolution by *∼* 30% for both GCaMP6f and GCaMP5k, a gain which can be considered as very significant given that it is here obtained with no prior knowledge of the convolution kernel. A more modest gain is obtained for GCaMP6s, likely due to the higher noise and larger rise time associated with this dataset.

### B. Light-sheet Imaging of Zebrafish

Compared to standard fluorescence microscopy technique, such as confocal or two-photon point-scanning technique, light-sheet imaging allows for a parallelization of the recording, yielding ~ 100-fold increase in data-throughput [19]. When applied to zebrafish larvae, this enables simultaneous recording of the quasi-entirety of the neurons (*∼* 100, 000 units) at typically 1 brain/second. The BSD algorithm might prove to be particularly useful for such experiments, as the size of individual datasets precludes supervision. Furthermore, the gain in speed with respect to con-oopsi should also be beneficial as it may allow one to carry out the spike inference on the fly.

To illustrate this latter claim, we test con-oopsi and BSD inference algorithms on a typical whole-brain recording, consisting of 1,800 successive volumetric stacks sampled at 1 stack/second, each of them comprising 20 z-sections. The experiment is performed on a 5 dpf larva expressing the GCaMP5 reporter panneurally. Voxels were clustered by a factor X. Hence, 255463 fluorescence traces encompassing the brain volume are processed independently. The baseline is computed as described before and the spike deconvolution is then carried out using both BSD and con-oopsi on an Intel Xeon Phi (28 cores) computer. In line with our observations of Section 2D2, we find that BSD achieves a 7-fold increase in speed compared to con-oopsi, see table V. Under these experimental conditions, the computation time with BSD match the duration of the experiment itself (20 minutes), making possible real-time spike inference. Importantly, the computation time per voxel is fairly stable with BSD, whereas some voxels use up to 200 times more time to be processed than others with con-oopsi.

These brain-scale simultaneous recordings allow one to compute the correlation of neuronal pairs activity, which might then be used to extract information regarding the large-scale functional organization of the brain. In this context, we examine whether the correlation statistics of the spike-inferred signals may be significantly different from the one computed using the raw DFF signals. For this purpose, we use a 2D recording acquired at 20 frame/second for 20 minutes in a 5dpf-old zebrafish larva expressing the genetically encoded indicator GCaMP3 (elavl3:GCaMP3). Automatic segmentation allowed us to identify 8082 individual neurons or neuropil regions of similar area, and the inference is then carried out on the ROI-averaged fluorescence traces. The rise and decay times are inferred for all neurons (see supplementary Figure). The average values of these two time-constants are then used to perform spike inference.

Figure 7a displays the time-averaged image of the brain section. Fluorescent traces and associated inferred spike trains for 5 representative neurons located in various brain regions are shown in Figure 7b. As expected, the deconvolved spike trace appear much sparser and less noisy than the original fluorescent signal. The pair-wise correlations, corrected for uniform coherent noise, are then computed for both the raw DFF signal and the inferred spike traces. We find the correlation distribution to be much more peaked after deconvolution (Figure 7c) which reflects in the more uniform appearance of the associated correlation matrix (Figure 7d).

**FIG. 7:**
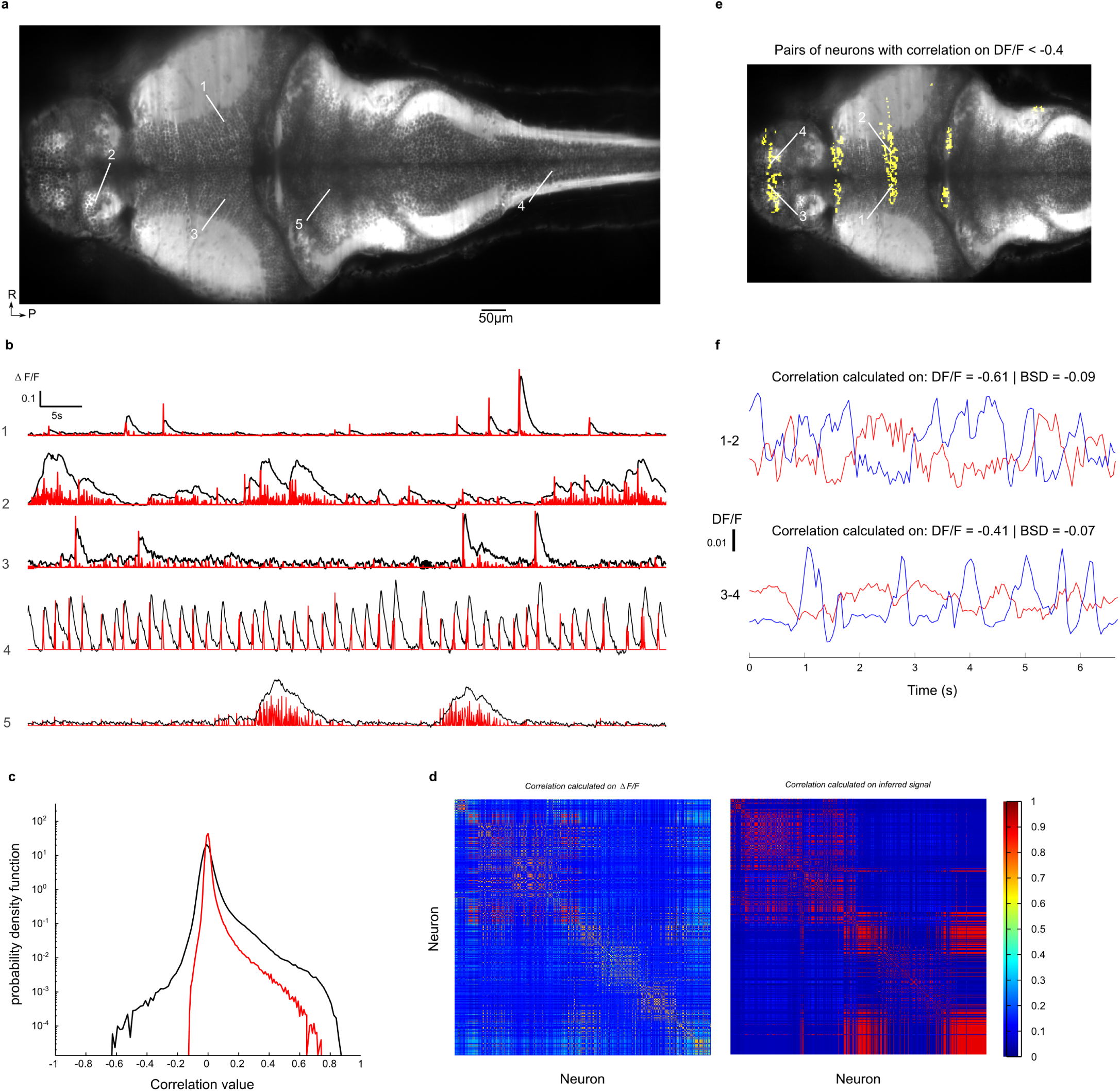
(a) Bottom: Individual traces of 5 neurons recorded at 20Hz from 6dpf larvae, in black curve DF/F, in red resulting signal from BSD deconvolution algorithm. Top: Time-averaged image of a brain slice of the larva, the white arrows give the location of the 5 neurons. (b) Distribution of pair-wise correlations of DF/F (black) and signal after BSD deconvolution. The data were obtained from a 20Hz, 20 min long experiment on a 6dpf larva. (c) Time-averaged image of a brain slice of the larva. In yellow, pairs of neurons that display a correlation on DF/F inferior to -0.4. Top: Pair of neuron DF/F traces that display a pair-wise correlation calculated on DF/F of -0.61 and a pair-wise correlation calculated after BSD deconvolution of -0.09. Down: Pair of neuron DF/F traces that display a pair-wise correlation calculated on DF/F of -0.41 and a pair-wise correlation calculated after BSD deconvolution of -0.07. (e) Correlation matrix computed from DF/F. (f) Correlation matrix computed from the signal after deconvolution

**FIG. 8:**
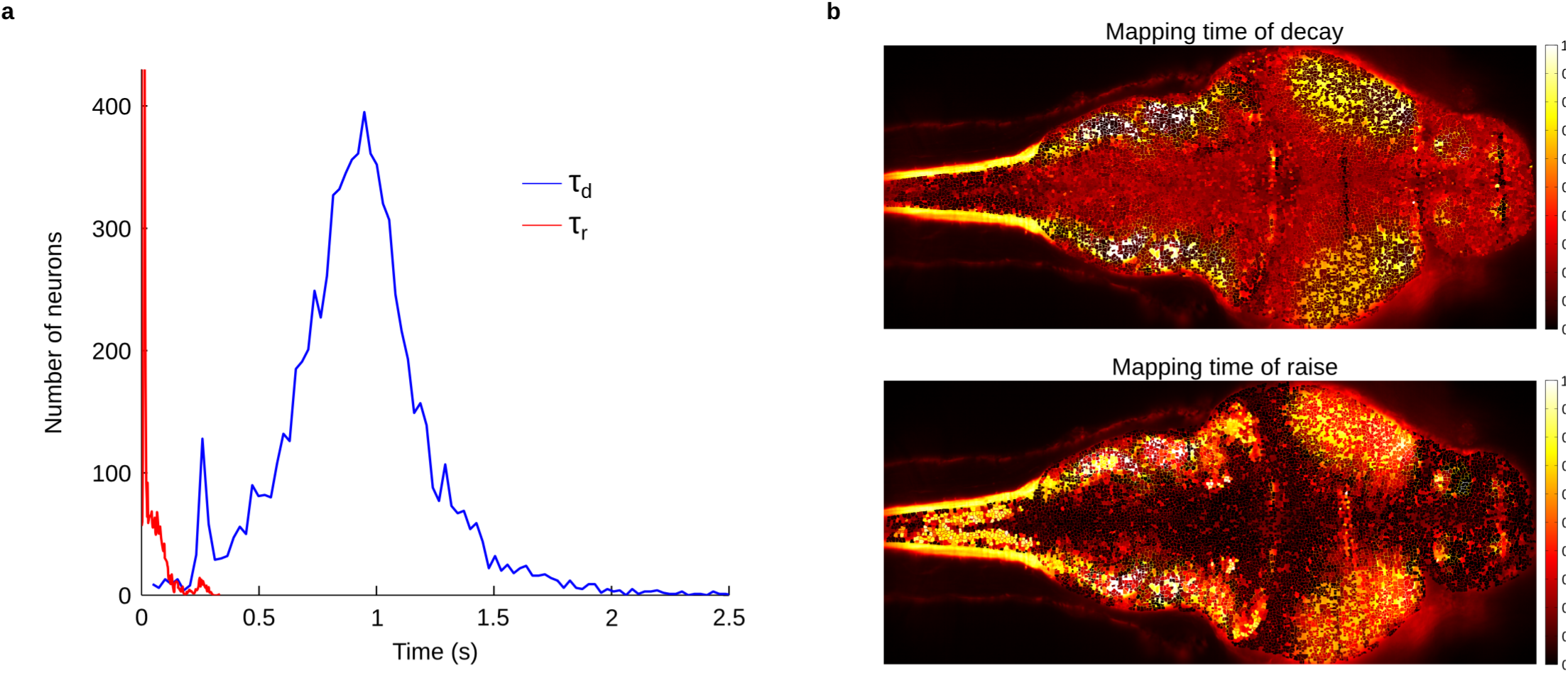
(a) Distribution of rise and decay time. (b) Mapping of rise and decay time across a neurons

This difference may have two possible origins. First, it may reflect the gain in temporal precision brought along by the spike inference, which may reduce the correlation of neuronal pairs that tend to discharge coherently (due to common inputs for instance), but with a slight systematic time-lag. A second explanation is related to the denoising property of the inference. In light-sheet imaging, the noise tends to display significant spatial correlation. This is notably due to the motion of small absorbing objects such as red cells that project elongated shadows and produce characteristic streaking features. Provided that these artifacts have characteristic timescales distinct from the spike-induced fluorescent transient, they are not interpreted as actual spike by BSD. This latter interpretation is confirmed by the fact that the highly negatively correlated pairs in the raw fluorescence signals are mostly confined within thin bands aligned along the beam direction (Figure 7e). For the same neuronal pairs, the correlation value computed from the inferred signal is thus largely reduced (Figure 7f).

## Conclusion

One may expect that spike inference algorithms become an important tool for functional imaging experiments, as they allow one to recover a signal closer to the underlying spiking activity with reduced noise and better temporal resolution. However, current implementations suffer from various limitations, including long computational time, the need for arbitrary setting model parameters, the sensitivity of the outcome to experimental conditions and, more generally, a paucity of theoretical understanding of the expected performances. These drawbacks have hampered the generalization of these algorithmic methods, such that their use remain in practice relatively limited among neuroscientists.

Here we introduced a novel non-negative sparse algorithm, named Blind Sparse Deconvolution, which was designed to specifically address these issues. Compared to previous works, this fully unsupervised algorithm features higher computational speed, accuracy and adaptability. Moreover, it incorporates theoretically-grounded framework to derive estimates of the expected deconvolution performance in terms of temporal accuracy and precision-recall of the inferred spike train. These information may be used before recording as guidelines for experimental design, or a posteriori to estimate error rates. BSD also features temporal super-resolution, which we showed to significantly increase the temporal resolution on real data. As both calcium reporters and imaging methods will gain in sensitivity and speed, this capability may provide sufficient temporal precision to investigate, using calcium imaging, the role of spike-timing in extended networks dynamics. This package stands as a complement to libraries that address other challenges of calcium imaging processing, such as the spatial filtering.

## Annex A: Stability of the single spike solution and the half-spike problem

In Section 2, we have not studied the stability of the single-spike solution. We study it here, and discuss when it is a global optimum. Assuming 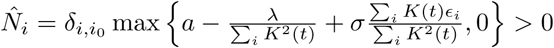 and looking for the stability of the solution, w.r.t the other coordinates, we find:

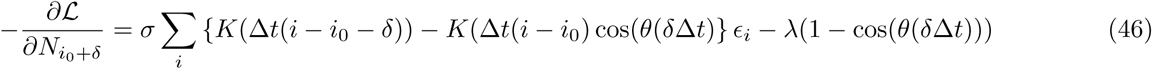

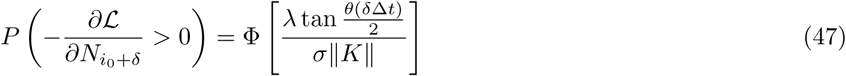

Where 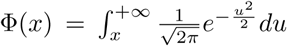. Therefore, the Dirac solution is stable only if the above probability is small enough for all values of *δ*. Far away from the spike *δ → ∞*, the angle 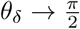 and we recover 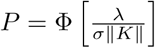, as in the spikeless signal. On the other hand, the smaller *δ*, the smaller *θ*_*δ*_ and the probability is higher. For *λ* = *λ*_*BSD*_ and low noise, the above probability reduces to 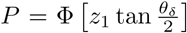; the Dirac solution can become unstable. In practice, the result depends on the level of noise: for low *σ*, the optimum remains close to the Dirac solution, whereas for high noise, we can find ‘half-spikes’ solutions, of the form 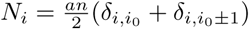

## Annex B: Kernel inference: proof of convergence and fast algorithm

We prove here that for isolated spikes and small noise, the cost function 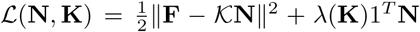 admits solution *K* = *K*^0^ as local minimum. Denoting 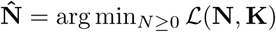

For a signal with a single spike 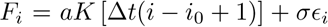, if the noise is small and *K* is close enough to *K*_0_, we have: 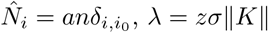 Optimizing over n yields:

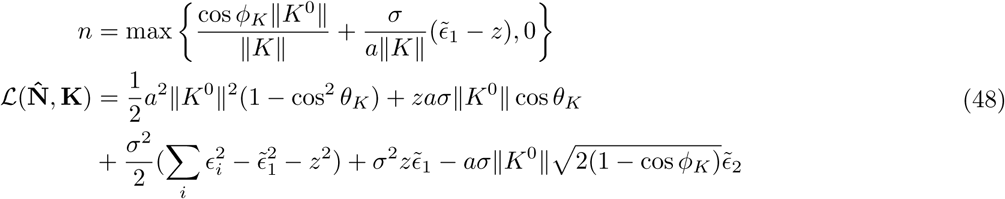

Where:

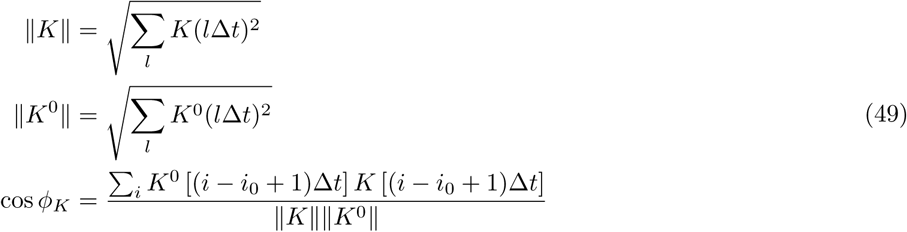

Note that we recover Eq 17 when *K* = *K*^0^. For a signal of multiple isolated spikes 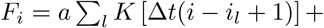, with 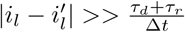, a similar solution 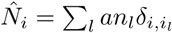 can be derived, and *L* is self averaging:

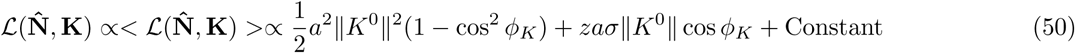

Hence, the function depends on **K** only through cos *θ*_*K*_. One can check that when 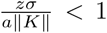, the minimum is reached at cos *φ*_*K*_ = 1, *i.e. K* = *K*^0^. This concludes the proof. Although we can not prove more about the radius of convergence, good convergence was achieved in practice after starting from the initialization.

In practice, the optimization with respect to *K* can be performed efficiently using standard temporal regression tricks. Observe that:

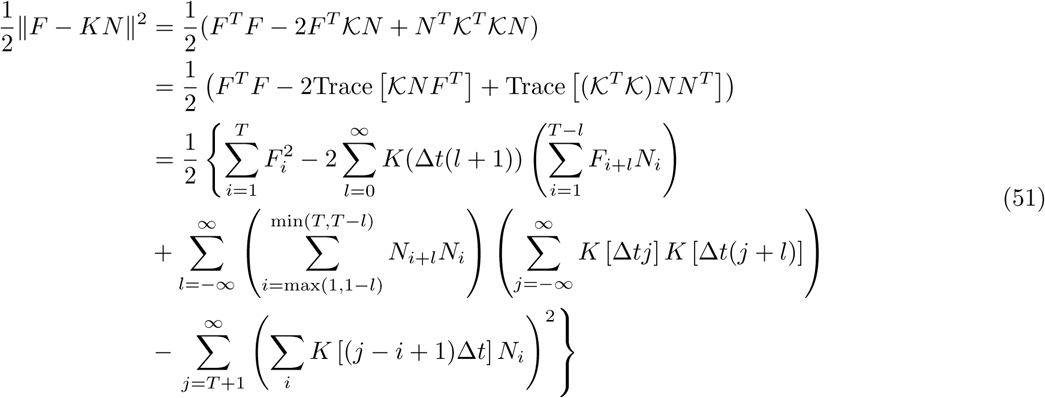

To go from the second line to the third line, we used the translation invariance property of *K*, the causality of *K* (*K*_*ij*_ = 0*∀j ≥ i*) and wrote 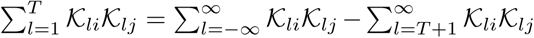 Hence, *L*(**N**, **K**) depends on *N* and *F* only through:

- the sums 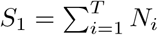 and 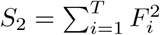
- the unnormalized cross-correlation between fluorescence and inferred spikes 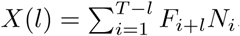.
- the unnormalized autocorrelation function of the inferred spikes 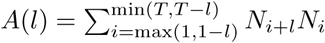
- the boundary term 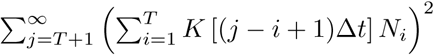

The first three terms can be precomputed in *𝒪*(*T*) once for all, and the second and third up to a cutoff 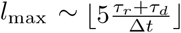, such that *K*(*l*_max_) *<<* 1. The last one can be computed in *𝒪*(*l*_max_), by noting that after *T*, the convolved spikes is a double exponential, with coefficients depending on the *∼ l*_max_ last time bins. Overall, the cost function can be evaluated in *𝒪*(*l*_max_) and optimized efficiently.

## Annex C: Detailed computations for the response function estimation

We assume a noisy single spike signal, *F*_*i*_ = *aK* [Δ*ti − t*_0_] + *σE*_*i*_, where we write formally *t*_0_ = Δ*t* (*i*_0_ *−* 1 + *r*_0_), with *r*_0_ *∈* [0, 1[; *i.e.* the spike is emitted before measurement *i*_0_. The likelihood27 becomes:

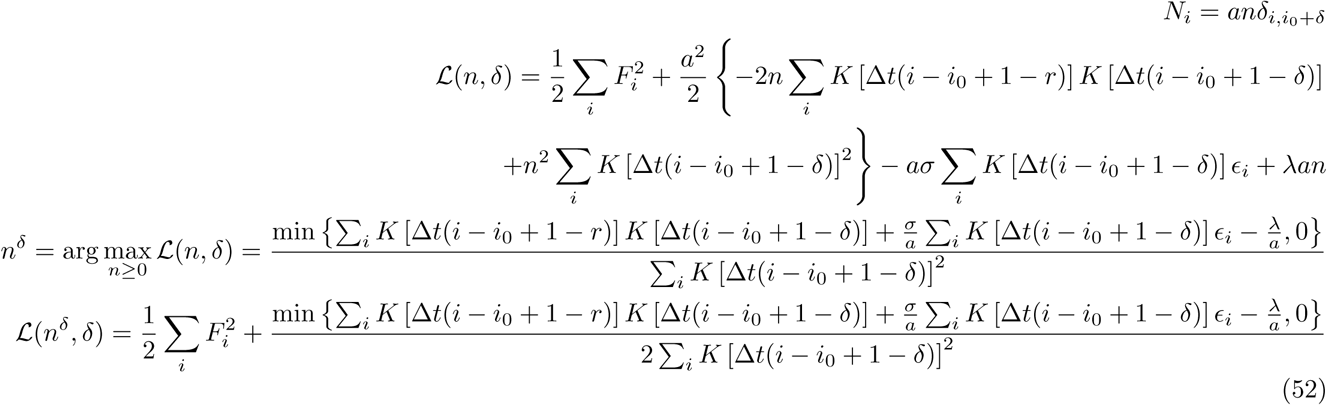

In the last expression, the term 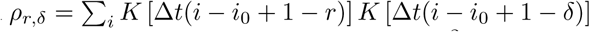 can be computed analytically for all *δ* and *r* and is independent of *i*_0_; the term 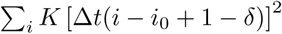 is the usual 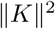 and the term involving noise can be rewritten by introducing new, correlated gaussian noises:

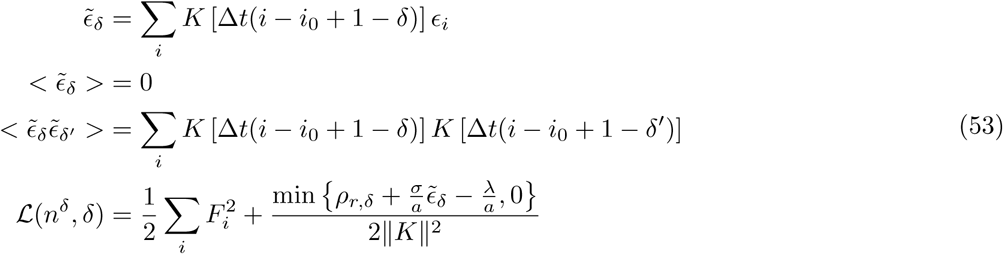

For a given *r* and noise realization, we can thus compute the optimal *δ* - and by Monte Carlo averaging, we obtain an estimate of the probability distribution *P* (*δ r*). To obtain a response function in continuous time, it is then transformed into a continuous piecewise-constant probability density through: 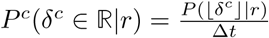.

And the overall response function is obtained by averaging over *r*, yielding:

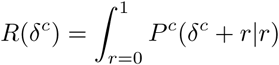

In practice, *R* and *r* are computed over a discrete grid of the form 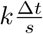.

For the super-resolution case, the computation is almost the same; the only difference being that we reconstruct the spikes with a thinner resolution.

## Annex D: Proof of unbiased estimation for super-resolution

We show here that the choice 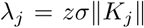 is best suited for an unbiased (in time) reconstruction of the spikes. We consider again the single-spike setting, with a single spike of *a* at position *k*_0_ = (*i*_0_ *−* 1)*s* + *r*_0_, for which 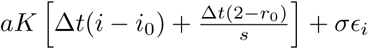

We now look for optima of *L*(**N**, **K**) of the form 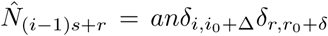 Note that instead of doing this computation, we can simply observe that it is a special case of Annex B, using reference kernel *K*^0^(*t*) *≡ K*(*t*), and measurement kernel 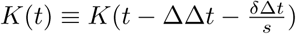.

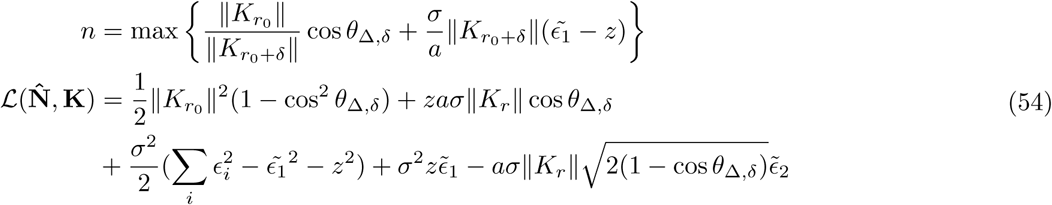

Where:

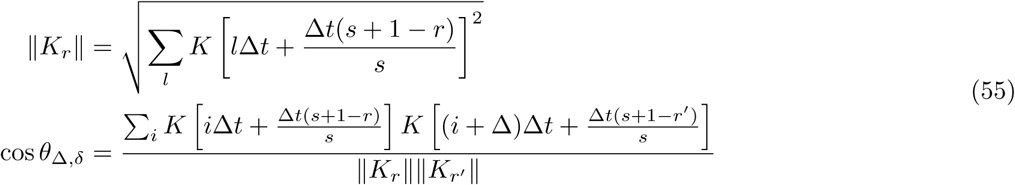

And 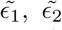 are gaussian noises of variance unity (see Annex B). Thus, the optimum over Δ, *δ* is with highest probability *δ* = Δ = 0, and the estimator is unbiased. Note that this result is expected: using the equivalence with a LASSO regression developed in Sec. 5, we know that the coefficients (here, the spikes) are correctly estimated with a uniform *λ* only when the features (Here, *K*) are normalized to unity 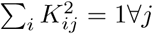

## Annex E: Kernel inference in the super-resolution setting

Since the convolution matrix *K* is not fully translation invariant in the super-resolution setting, the estimation of the kernel is slightly different. For the initial estimation, Eqn. 28 becomes:

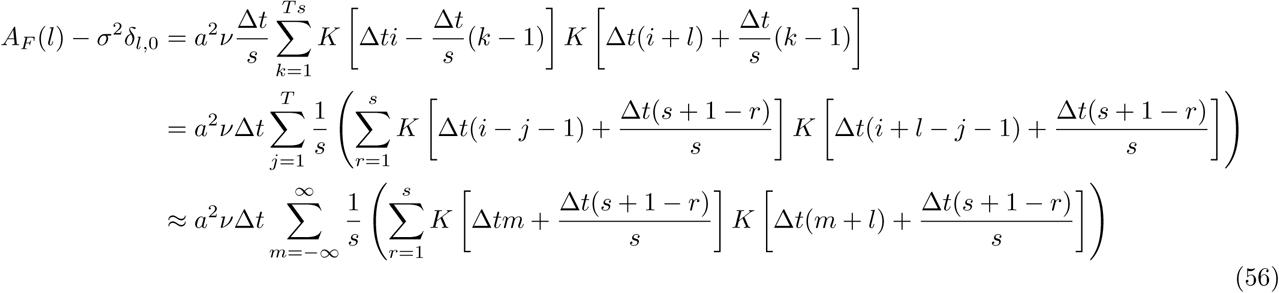

For *s >* 1, this formula is different from Eqn. 28. It can be shown (see Annex E) that the right-hand side has a well-defined limit when *s→ ∞*, ie in the continuous setting.

Similarly, the iterative kernel update Eqn. 51 is different:

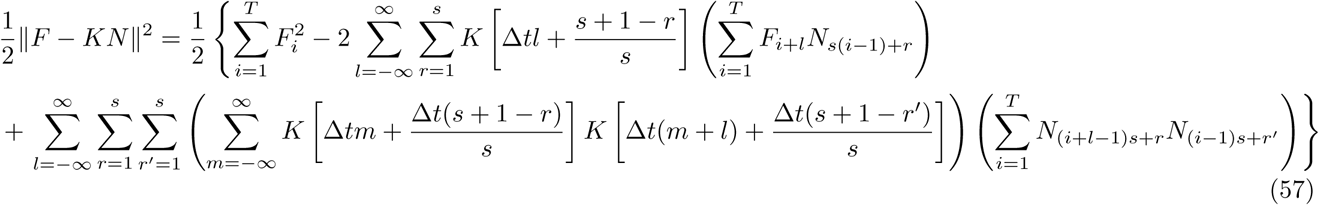

The sparsity penalty becomes:

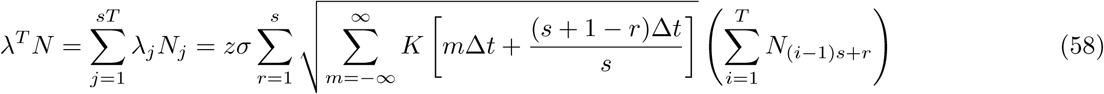

Hence, *L*(**N**, **K**) now depends on **F** and **N** through the following quantities:

- the sum 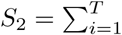
- the sums vector 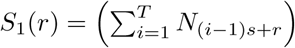
- the cross-correlation matrix 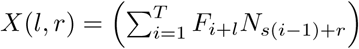
- the autocorrelation tensor 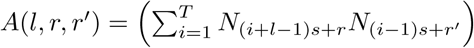

Altogether, the cost function can be evaluated relatively fast. Note that the complexity of the kernel optimization is now 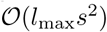.

## Annex F: Various explicit formulas for the double exponential kernel

Various useful formulas for blind sparse deconvolution are consigned, here for double exponential kernels.

**Kernel normalization**. We normalize *K* such that max_*t≥*0_ *K*(*t*) = 1. This gives:

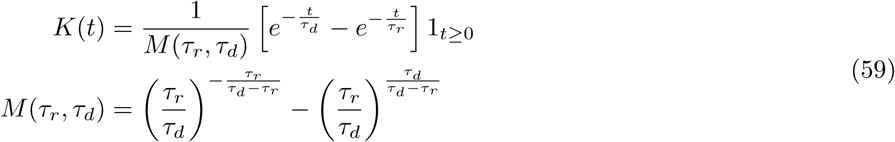

**Kernel norms**. The *L*_1_ and *L*_2_ norms are computed as follow:

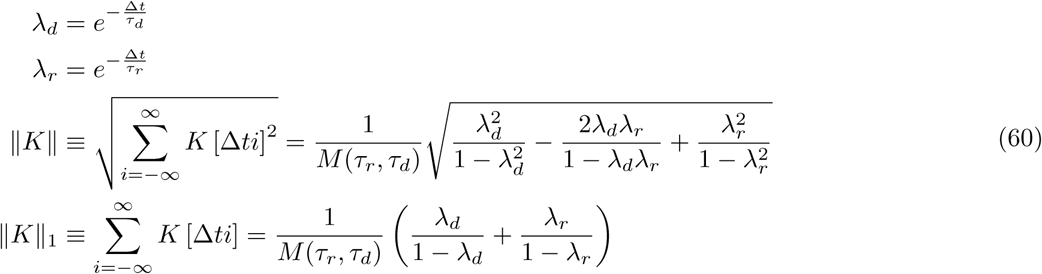

Kernel norms for super-resolution. The *L*_1_ and *L*_2_ norms for a spike emitted at time (*j −* 1)*s* + *r, r ∈* [1, *s*] are given by:

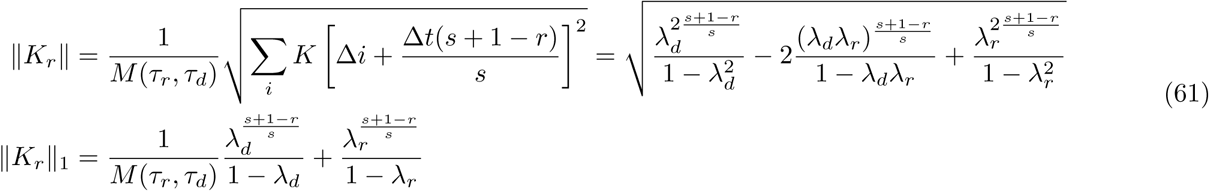

**Kernel overlaps** Useful for assessing temporal uncertainty and for kernel estimation

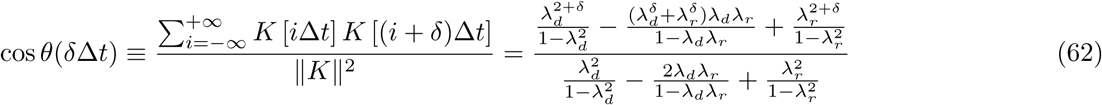

**Boundary term** The estimation of the kernel involves the computation of the following boundary term:

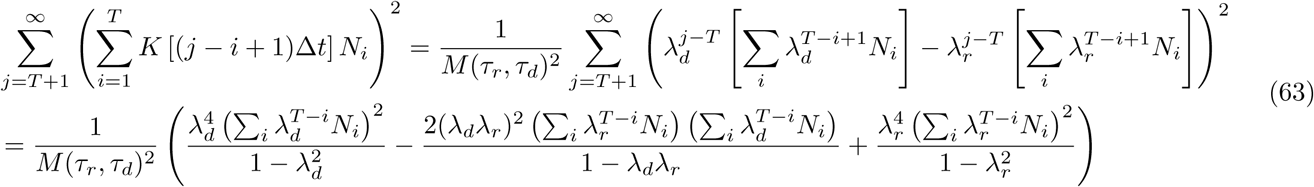

**Kernel overlaps for super-resolution** Useful for assessing temporal uncertainty and for kernel estimation

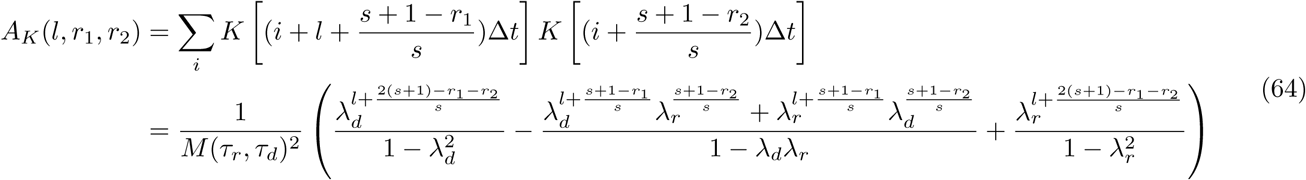

In particular:

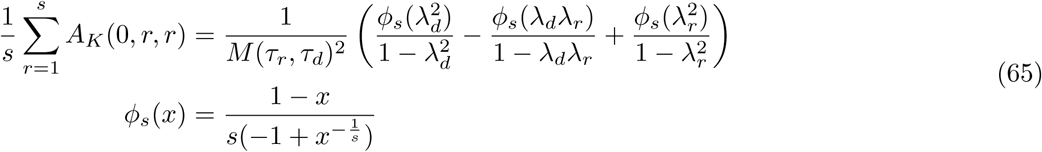

### Boundary-term for super-resolution

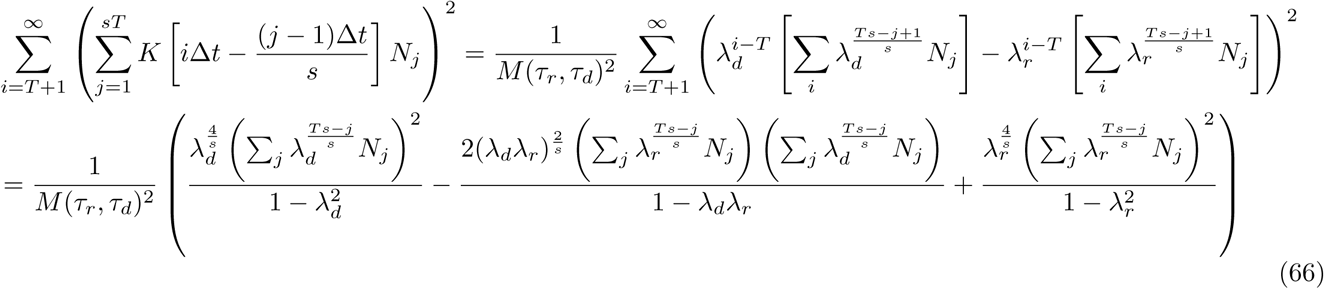

## Annex G: Heterogeneity in rise and decay time constants in Zebrafish

Application of BSD to zebrafish data yields heterogeneous distributions of rise and decay times. This means that different regions show different patterns of fluorescence bursts. We see that the heterogeneities have a spatial structure: in particular neurons in X tend to have longer time constants, whereas neurons in have shorter time constants.The two possible explanations are that the spike patterns are different in these regions (*e.g.*, regular vs sparse spike trains), and/or that the expression of GCaMP is significatively different. Overall, they motivate the use of heterogeneous time constants.

**TABLE I:** Comparison of BSD, fast-oopsi and con-oopsi computational speed. For con-oopsi, under-estimation of the noise level *σ*, even for synthetic data, can lead to a large increase in computational time

| $N_{frames}$ | BSD/ fast-oopsi (s) | con-oopsi (s) | con-oopsi $\sigma$ user-provided |
| --- | --- | --- | --- |
| $10^4$ | 0.7 | 2.4 | 2.1 |
| $5 \cdot 10^4$ | 2.6 | 8.2 | 8.3 |
| $2 \cdot 10^5$ | 10 | 84 | 45 |
| $5 \cdot 10^5$ | 27 | 529 | 128 |
| $10^6$ | 49 | 1594 | 290 |

**TABLE II:** Data sets summary

|  | GCaMP5k | GCaMP6s | GCaMP6f |
| --- | --- | --- | --- |
| Number of recordings | 9 | 21 | 37 |
| Total number of spikes | 2735 | 2103 | 4535 |
| Average $\tau_r$ | 0.058 | 0.078 | 0.024 |
| Average $\tau_d$ | 0.52 | 0.88 | 0.38 |
| Median Noise level $\frac{\sigma}{a}$ | 0.35 | 0.40 | 0.30 |

**TABLE III:**
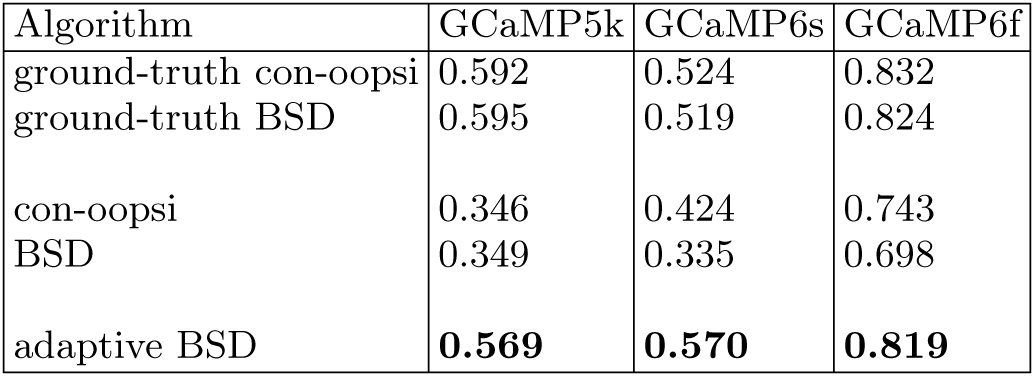
Spike detection performance (AUC)

**TABLE IV:**
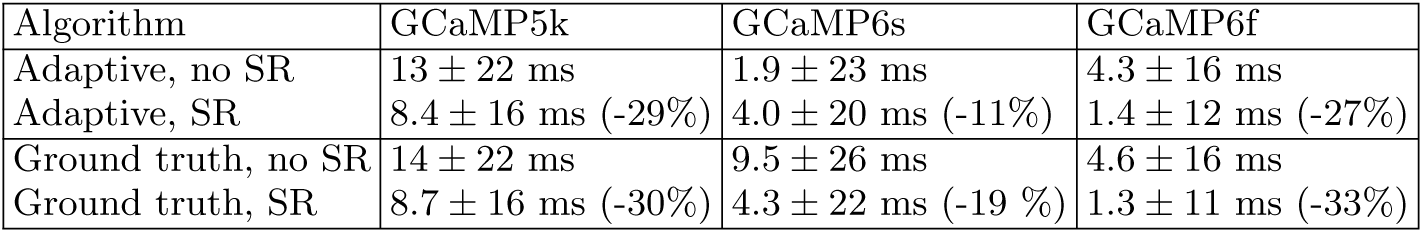
Temporal accuracy of various algorithms: offset *μ* and width *σ* of the response function (ms). The relative gain in root square distance 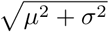 is displayed. The frame rate interval is 16.6ms

| Algorithm | GCaMP5k | GCaMP6s | GCaMP6f |
| --- | --- | --- | --- |
| Adaptive, no SR | $13 \pm 22$ ms | $1.9 \pm 23$ ms | $4.3 \pm 16$ ms |
| Adaptive, SR | $8.4 \pm 16$ ms (-29%) | $4.0 \pm 20$ ms (-11%) | $1.4 \pm 12$ ms (-27%) |
| Ground truth, no SR | $14 \pm 22$ ms | $9.5 \pm 26$ ms | $4.6 \pm 16$ ms |
| Ground truth, SR | $8.7 \pm 16$ ms (-30%) | $4.3 \pm 22$ ms (-19 %) | $1.3 \pm 11$ ms (-33%) |

**TABLE I:**
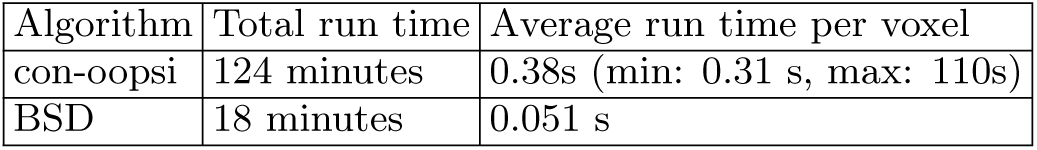
Time for perfoming deconvolution on voxelated data

